# Tgif1-deficiency impairs cytoskeletal architecture in osteoblasts by activating PAK3 signaling

**DOI:** 10.1101/2023.11.04.565288

**Authors:** Simona Bolamperti, Hiroaki Saito, Sarah Heerdmann, Eric Hesse, Hanna Taipaleenmäki

**Affiliations:** Molecular Skeletal Biology Laboratory, Department of Trauma, Hand and Reconstructive Surgery, University Medical Center Hamburg-Eppendorf, Martinistraße 52, 20246 Hamburg, Germany; Institute of Musculoskeletal Medicine, LMU University Hospital, LMU Munich, Fraunhoferstraße 20, 82152 Planegg-Martinsried, Germany; Musculoskeletal University Center Munich, LMU University Hospital, LMU Munich, Fraunhoferstraße 20, 82152 Planegg-Martinsried, Germany

**Keywords:** osteoblast spreading, Tgif1, PTH, bone regeneration, PAK3

## Abstract

Osteoblast adherence to bone surfaces is important for remodeling of the bone tissue. This study demonstrates that deficiency of TG-interacting factor 1 (Tgif1) in osteoblasts results in altered cell morphology, reduced adherence to collagen type I-coated surfaces, and impaired migration capacity. Tgif1 is essential for osteoblasts to adapt a regular cell morphology and to efficiently adhere and migrate on collagen type I-rich matrices *in vitro*. Furthermore, Tgif1 acts as transcriptional repressor of p21-activated kinase 3 (PAK3), an important regulator of focal adhesion formation and osteoblast spreading. Absence of Tgif1 leads to increased PAK3 expression, which impairs osteoblast spreading. Additionally, Tgif1 is implicated in osteoblast recruitment and activation of bone surfaces in the context of bone regeneration and in response to parathyroid hormone 1-34 (PTH 1-34) treatment *in vivo*. These findings provide important novel insights in the regulation of the cytoskeletal architecture of osteoblasts.

## Introduction

Bone remodeling is a highly coordinated and continuously ongoing process involving osteoblasts, osteoclasts, and osteocytes, that interact with each other and function in concert to maintain bone mass and preserve the integrity of the skeletal system (Baron and Hesse, 2012). This process is essential for bone development, growth, maintenance, and repair as well as for the functionality of pharmacological interventions (Kenkre and Bassett, 2018). Osteoblasts, derived from mesenchymal stem cells, play a pivotal role in synthesizing and depositing collagen type I-rich bone matrix (Ponzetti and Rucci, 2021). To achieve this, osteoblasts undergo changes in cell shape and migrate on bone surfaces to lay down matrix at places where new bone needs to be formed, necessitating the regulation of cell movement and shape acquisition (Jones and Boyde, 1977; Eleniste et al., 2014; Thiel et al., 2018).

Migration of osteoblasts involves movement from their origin to specific locations within the bone tissue and occurs during both embryonic development and adult bone remodeling. Multiple factors influence osteoblast migration, including growth factors, cytokines, signaling pathways like bone morphogenetic proteins (BMPs) and transforming growth factor-beta (TGF-β) as well as interactions with other cells and the extracellular matrix (Thiel et al., 2018). These signals not only promote migration but also regulate the expression of adhesion molecules, cytoskeletal rearrangements, and cellular contractility.

Osteoblasts exhibit various morphological shapes based on their differentiation stage and the local microenvironment, ranging from elongated spindle-like morphology to cuboidal appearance (Tsuji et al., 2022). The cytoskeleton, primarily composed of actin filaments, microtubules, and intermediate filaments, plays a central role in determining cell shape. Actin filaments, among other cytoskeletal elements, are involved in cell polarization, protrusion formation, and contractility during migration (Schaks et al., 2019; Tang and Gerlach, 2017; Etienne-Manneville, 2004). Signaling molecules, including Rho GTPases such as RhoA, Rac1, and Cdc42, regulate the cytoskeleton by controlling actin polymerization and organization, contributing to the formation of structures like lamellipodia, filopodia, and stress fibers (Schaks et al., 2019). These structures are essential for cell migration, enabling attachment to the extracellular matrix and generating the necessary force for movement. Integrins, transmembrane proteins mediating cell-extracellular matrix interactions, also play a critical role in osteoblast migration and cell shape regulation. They anchor cells to the matrix, transmit signals regulating migration and cytoskeletal dynamics, and activate intracellular pathways affecting cell shape, migration speed, and adhesion strength (Kechagia et al., 2019; SenGupta et al., 2021; Hood and Cheresh, 2002; Thiel et al., 2018). Thus, osteoblast migration and cell shape acquisition are interconnected processes crucial for bone development, growth, remodeling, and repair. Further investigation of the mechanisms underlying osteoblast migration and cell shape regulation is therefore important to better understand bone mass maintenance, treatment of bone-related disorders, and bone repair.

Fractures occur in the context of high-energy trauma but also due to bone fragility and represent a severe damage of bone integrity, requiring subsequent tissue regeneration. In response to fracture, inflammatory cells including neutrophils and macrophages are recruited to the site, releasing signaling molecules such as cytokines and growth factors. These molecules create a favorable environment for osteoblast migration and bone formation (Einhorn and Gerstenfeld, 2015). Osteoblast precursors derived from mesenchymal stem cells, or the periosteum migrate alongside sprouting vessels towards the fracture gap and differentiate into mature bone-forming osteoblasts (Maes et al., 2010). Once osteoblasts have reached the fracture site, production and deposition of new bone matrix is initiated to facilitate fracture repair (Maes et al., 2010; Einhorn and Gerstenfeld, 2015). During this process, chemotactic signals released by surrounding tissues and cells, including platelet-derived growth factor (PDGF), transforming growth factor-beta (TGF-β), and bone morphogenetic proteins (BMPs) guide osteoblast migration (Dirckx et al., 2013; Einhorn and Gerstenfeld, 2015; Thiel et al., 2018). Furthermore, migration of osteoblasts is facilitated by the dynamic rearrangement of the cytoskeleton, primarily composed of actin filaments (Thiel et al., 2018). Structures like lamellipodia and filopodia, enriched with actin, enable osteoblasts to extend and protrude in the direction of migration, interacting with the extracellular matrix and providing necessary traction (Casati et al., 2015; Jafari et al., 2019).

Movement of osteoblasts is an important but often underestimated component of the pharmacologically induced gain in bone mass in response to the treatment with bone anabolic drugs. Teriparatide, a recombinant form of the first 34 amino acids of human parathyroid hormone (rhPTH1-34; hereafter PTH), is a bone anabolic drug used to treat severe osteoporosis, characterized by low bone mineral density and an increased fracture risk (Neer et al., 2001; Black and Rosen, 2016). PTH treatment stimulates bone formation by increasing the number, differentiation, and activity of osteoblasts (Pettway et al., 2008; Silva et al., 2011; Ogita et al., 2008). In addition, PTH promotes the recruitment of osteoblast precursor cells and activates quiescent lining cells on bone surfaces, transforming them into active matrix-forming osteoblasts (Nishida et al., 1994; Dobnig and Turner, 1995; Kim et al., 2012). Furthermore, PTH increases the migration of mesenchymal precursor cells (Lv et al., 2020) and augments bone remodeling (Saito et al., 2019), which attracts osteoblasts (Dirckx et al., 2013). During osteoblast migration, cytoskeletal dynamics promotes actin filament remodeling and the formation of filopodia and lamellipodia (Lomri and Marie, 1990). These structures are essential for cell attachment to the extracellular matrix during migration. By enhancing cytoskeletal dynamics, PTH facilitates osteoblast movement (Thiel et al., 2018). PTH also increases the expression and activation of integrins, which mediate interactions between cells and the extracellular matrix (Gronthos et al., 2001; Kaiser et al., 2001). These combined pharmacological effects of PTH treatment result in an increase in bone formation and bone mineral density, which improves bone strength and ultimately reduces fracture risk (Neer et al., 2001; Black and Rosen, 2016).

Recently, we reported the important role of the homeodomain protein TG-interacting factor 1 (Tgif1) in osteoblast differentiation, activity, and bone formation (Saito et al., 2019). Furthermore, Tgif1 was identified as PTH target gene, and in the absence of Tgif1, PTH treatment failed to increase bone mass in mice (Saito et al., 2019). These findings demonstrate the importance of Tgif1 as novel regulator of bone remodeling and emphasizes its essential involvement in mediating the full bone anabolic effect of PTH treatment. Building upon these observations, we investigated the role of Tgif1 in osteoblast morphology, adherence, and migration. Our findings demonstrate that Tgif1-deficient osteoblasts display an altered morphology, reduced adherence to collagen type I-coated surfaces, impaired migration capacity and decreased spreading compared to control cells. These deficits are associated with a compromised formation of focal adhesions and an upregulated expression of p21-activated kinase 3 (PAK3), which we further investigated due to its implication in cell migration and adhesion (Liu et al., 2010). Mechanistically, Tgif1 exerts control over PAK3 expression through transcriptional repression. Thus, elevated PAK3 levels contribute to the observed defects in osteoblast spreading of Tgif1-deficient cells. By using translational approaches, we demonstrate that in the absence of Tgif1, the activation of bone surfaces by osteoblasts in response to bone repair and PTH treatment is diminished. Furthermore, we uncovered that PTH facilitates osteoblast spreading via Tgif1-PAK3 signaling.

Collectively, these findings increase the knowledge on processes governing osteoblast morphology, adherence, and migration. These novel insights are important to better understand mechanisms underlying bone remodeling and repair as well as the pharmacological effects of PTH treatment.

## Results

### Tgif1-deficiency alters osteoblast morphology, adherence and migration

Recently, we reported that in Tgif1-deficient mice the number and activity of osteoblasts are reduced, which contributes to a low turnover bone remodeling (Saito et al., 2019). Further histological examination revealed that bone surfaces were scarcely occupied by small and flat osteoblasts in mice bearing a germline deletion of Tgif1 (Fig. 1A) or a deletion of Tgif1 targeted to osteoblasts (Fig. 1B) compared to control littermates. This finding suggested that Tgif1-deficient osteoblasts might, in addition to an impaired bone matrix-producing capacity (Saito et al., 2019) be compromised in acquiring a physiological osteoblast morphology. This deficit could contribute to the low turnover bone phenotype of Tgif1-deficient mice.

**Figure 1.**
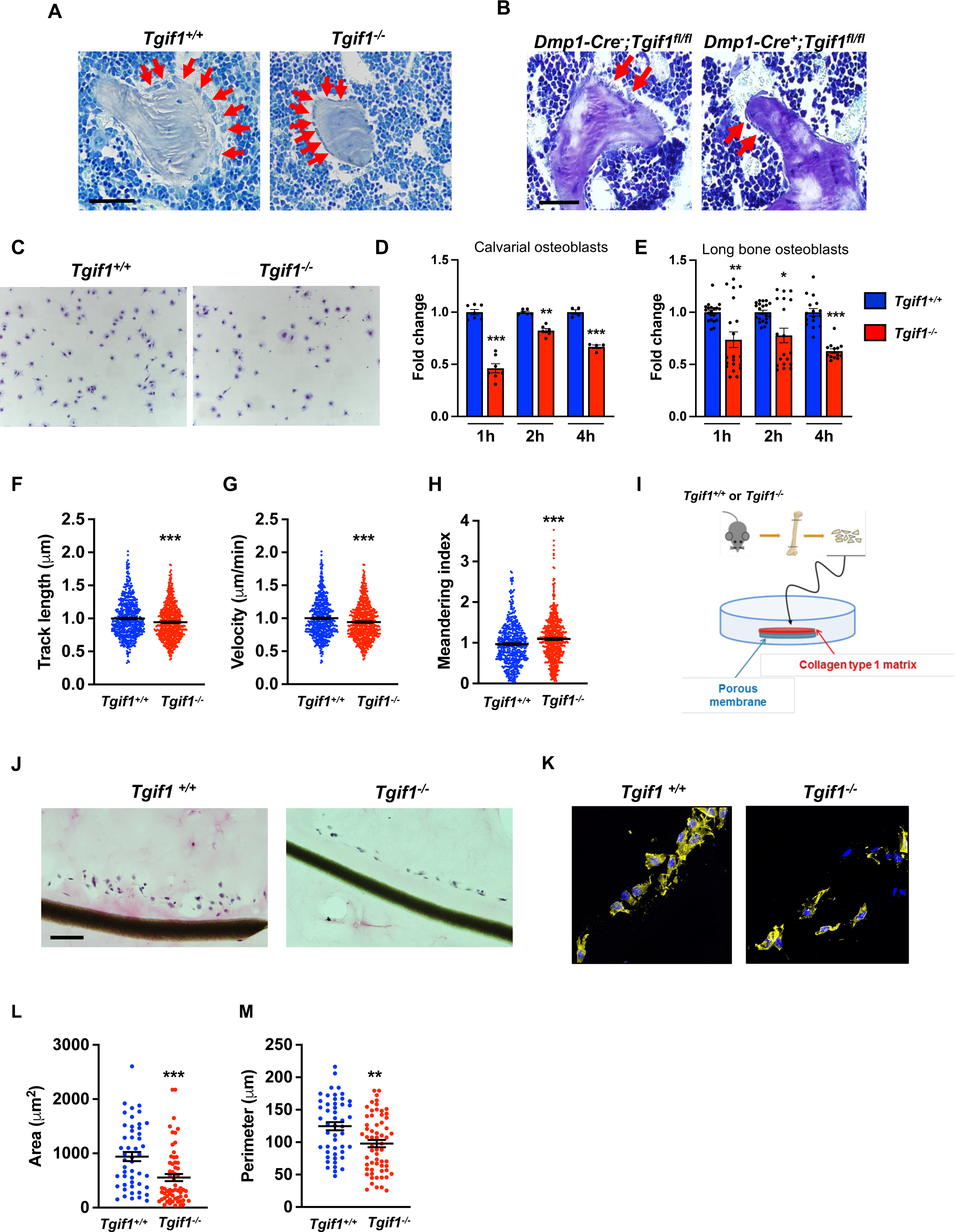
Loss of Tgif1 reduces osteoblast size *in vivo* and impairs osteoblast adhesion and migration *in vitro*. **(A)** Representative images of the femora from 12-week-old *Tgif1^+/+^* (n=4) and *Tgif1^-/**-**^* (n=4) mice stained with Toluidine blue. **(B)** Femora of 12-week-old *Dmp1-Cre^-^;Tgif1^fl/fl^*and *Dmp1-Cre^+^;Tgif1^fl/fl^* mice stained with toluidine blue. **(A, B)** Arrows indicate osteoblasts. Scale bars indicate 100 µM. **(C, D)** Osteoblasts isolated from calvariae of neonatal *Tgif1^+/+^*and *Tgif1^-/**-**^* mice upon adherence on Col-I coated surfaces for 1, 2 and 4 hours, fixation and staining with toluidine blue. **(C)** Representative images of Toluide blue-stained cells after 2 hours of adhesion. **(D)** Quantification of adherent *Tgif1^+/+^*and *Tgif1^-/**-**^* calvarial osteoblasts after 1, 2, 4 hours of adhesion on Col-I-coated surfaces. **(E)** Osteoblasts isolated from long bones of 8-week-old *Tgif1^+/+^* and *Tgif1^-/**-**^* mice upon adherence on Col-I coated surfaces for 1, 2 and 4 hours, fixation and staining with toluidine blue. Quantification of adherent cells at indicated time points. **(F-H)** Migration of calvarial osteoblasts obtained from neonatal *Tgif1^+/+^* and *Tgif1^-/**-**^* mice was analyzed using live cell imaging. Quantification of **(F)** track length, **(G)** migration velocity and **(H)** meandering index of *Tgif1^+/+^* and *Tgif1^-/-^* calvarial osteoblasts. **(I)** Long bones were harvested from *Tgif1^+/+^* and *Tgif1^-/**-**^* mice. Osteoblasts were isolated from bone chips and spread on Col-I matrices placed on porous membranes for 48h. Membranes were frozen, cut and stained with **(J)** H&E or **(K)** phalloidin (yellow) and DAPI (blue). Quantification of cell area **(L)** and cell perimeter **(M)** of *Tgif1^+/+^* and *Tgif1^-/-^*long bone osteoblasts on Col-I matrices. Scale bars indicate 50 µm. n = minimum of 3 independent experiments with technical duplicates. Unpaired t-test, *p<0.05, **p<0.01, ***p<0.001 vs. *Tgif1^+/+^*.

Inactive and early-stage osteoblasts are rather small and flat in their morphological appearance but enlarge and adopt a more cuboidal shape upon activation. Active osteoblasts produce extracellular matrix and have the capacity to migrate to sites of bone remodeling and repair (Dirckx et al., 2013). These features require the ability to adhere to surfaces and to change cell morphology to facilitate cell migration. In support of our hypothesis that Tgif1-deficiency compromises morphological plasticity of osteoblasts, calvarial- and long bone-derived osteoblasts obtained from Tgif1-deficient mice were impaired in their ability to attach to collagen type I-coated surfaces throughout a 4-hour time-course (Fig. 1C-E). To determine the ability of Tgif1-deficient osteoblasts to migrate, we performed live cell video microscopy of osteoblasts seeded on collagen type I-coated surfaces (Dang and Gautreau, 2018). This analysis revealed that osteoblasts lacking Tgif1 were impaired in their migration capacity reflected by a shorter track length, a reduced velocity and a less straight and therefore more meandering track path (Fig. 1 F-H, Fig. S1A). Thus, these *in vivo* and *in vitro* data indicate that Tgif1 is important for osteoblasts to adhere and migrate on collagen type I-rich surfaces. To confirm these findings in a 3-dimensional functional context, we obtained long bone osteoblasts from Tgif1-defcient mice and control littermates and seeded them on a collagen type I matrix placed over a porous membrane (Fig. 1I). In support of the previous observations, osteoblasts lacking Tgif1 adhered less to collagen type I matrix and were impaired in their capacities to migrate into the matrix compared to control cells (Fig. 1J). The experiment also confirmed that Tgif1-deficient osteoblasts were rather flat in morphology and smaller in size as determined by reduced cell area and perimeter (Fig. 1K-M). Collectively, these observations demonstrate that Tgif1 is indispensable for osteoblasts to adapt a regular cell morphology and to adhere and migrate on collagen type I-rich matrices *in vitro* and on bone surfaces *in vivo*.

### Tgif1-deficient osteoblasts are impaired in spreading and in forming focal adhesions

Adherence on osseous matrices and migration alongside bone surfaces towards sites of remodeling or repair are crucial functional features of osteoblasts (Dirckx et al., 2013; Thiel et al., 2018). Upon adherence and migration, osteoblasts undergo morphological changes and spread on bone surfaces, thereby decreasing their sphericity, a measure of how closely an object resembles the round shape of a sphere (Cruz-Matías et al., 2019). To determine the ability of Tgif1-deficient osteoblasts to spread, calvarial osteoblasts were isolated from *Tgif1^-/-^* mice and control littermates and subject to spreading on collagen type I-coated slides for 60 minutes in the presence of Calcein-AM, a dye that is fluorescent in living cells (Dejaeger et al., 2017). Compared to control cells, *Tgif1^-/-^*osteoblasts were significantly impaired in their ability to spread during the experiment (Fig. 2A, B).

**Figure 2.**
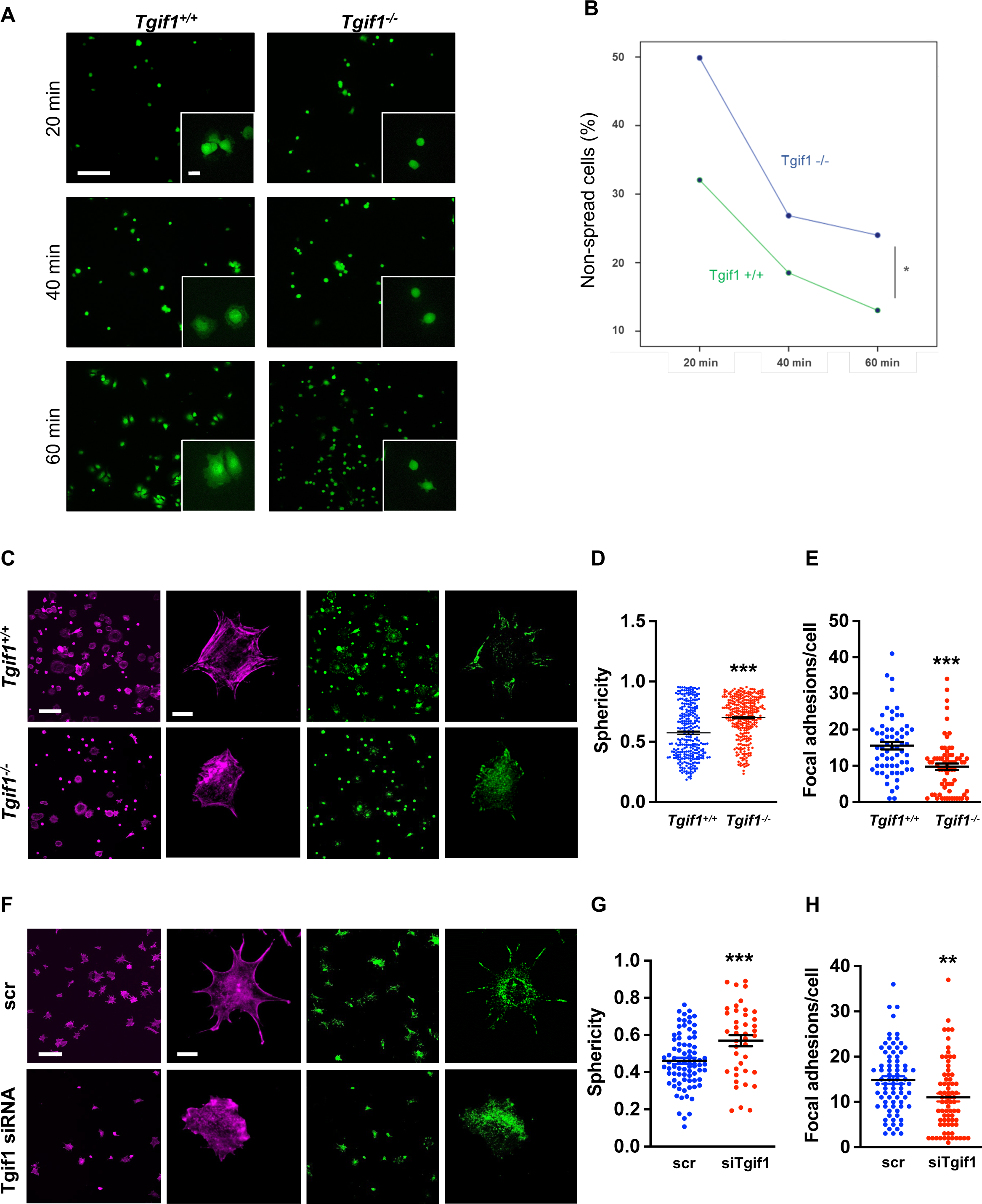
Tgif1-deficient osteoblasts are impaired to spread and form focal adhesions. **(A, B)** Calvarial osteoblast obtained from *Tgif1^+/+^* and *Tgif1^-/-^* mice adhered on Col-I coated slides for 20, 40 and 60 minutes. **(A)** Cell spreading was visualized by Calcein-AM. **(B)** The number of round (non-spread) cells was counted. Scale bar indicates 100 µm in low magnification images and 20 µm in the insets. n=6 independent experiments; Repeated Measures ANOVA, Estimated margins means test, *p<0.05 between genotypes. **(C)** *Tgif1^+/+^*and *Tgif1^-/-^* calvarial osteoblasts were allowed to adhere on Col-I coated slides for 60 minutes. Focal adhesion formation was visualized by paxillin staining (green) and actin cytoskeleton by phalloidin (magneta). Scale bars indicate 100 µm (lower magnification) or 10 µm (single cells). **(D)** Analysis of single cell sphericity after 3D reconstruction using IMARIS and **(E)** quantification of focal adhesions using Image J. (**F-H**) Tgif1 was silenced in OCY454 cells using siRNA and cells were allowed to adhere on Col-I coated slides for 60 minutes. **(F)** Focal adhesion formation was visualized by Paxillin staining (green) and cell protrusions by phalloidin (magenta). (**G**) Analysis of cell sphericity after 3D reconstruction and (**H**) quantification of focal adhesions. Unpaired t-test, \***\***\*p<0.001, **p<0.01 vs. *Tgif1^+/+^* (D, E) and scr (G, H).

Cell spreading is associated with the assembly and disassembly of actin filaments to form cell protrusions. Cell protrusions facilitate anchoring to the extracellular matrix via focal adhesion complexes (Cronin and Demali, 2021). To determine whether Tgif1 is implicated in actin filament- and focal adhesion assembly, *Tgif1^-/-^*and control osteoblasts were stained for phalloidin to visualize actin filaments and Paxillin, a major component of focal adhesions. Consistently, after 60 minutes of culture on collagen type I-coated surfaces, Tgif1-deficient osteoblasts were less spread and therefore had a greater sphericity (Fig. 2C, D). Furthermore, Tgif1-deficient osteoblasts that underwent spreading formed less cellular processes and focal adhesions compared to control cells (Fig. 2C, E).

To confirm the findings obtained from osteoblasts bearing a germline deletion of *Tgif1*, we transiently silenced *Tgif1* in cells of the OCY454 cell line, a cell model system resembling motile osteocytes and mature osteoblasts (Spatz et al., 2015), using siRNA (Fig. S1B). Consistent with the findings made by genetic deletion of *Tgif1*, siRNA-mediated silencing of Tgif1 in OCY454 cells resulted in an impaired cell spreading with a consecutive higher cell sphericity and a decrease in focal adhesion formation (Fig. 2F-H).

### PAK3 expression is increased in the absence of Tgif1 and impairs focal adhesion formation and osteoblast spreading

To investigate the molecular mechanisms underlying the impaired ability of *Tgif1^-/-^* osteoblasts to undergo morphological changes, expression and activation of the major focal adhesion components including FAK, Integrin β1, Src, talin, paxillin, p38 and LRG5 was quantified during cell spreading. However, no major changes were identified in the expression or activation of these molecules in the absence of Tgif1 (Fig. S2A).

Next, we examined the localization of Tgif1 to determine a possible interaction of Tgif1 with components of focal adhesions or actin filaments. Immunocytochemistry revealed that Tgif1 is predominantly localized in the nucleus and to some extend in the cytoplasm (Fig. S2B). However, cytoplasmic Tgif1 did not interact with focal adhesions, because no co-localization of eGFP-Tgif1 neither with paxillin nor with talin (Fig. S2B) was observed.

Given the predominantly nuclear localization of Tgif1 (Fig. S2B) and its established function as transcriptional repressor (Saito et al., 2019)(Wotton et al., 1999), we investigated whether absence of Tgif1 might alter the expression of genes involved in actin cytoskeletal assembly and re-arrangement as prerequisite of cell adhesion and migration. First, we quantified the expression of members of the cell division control protein 42 homolog (Cdc42), which are small GTPases of the Rho family that participate in the control of multiple cellular functions including cell migration and cell morphology (Hirsch et al., 2001). However, expression of none of the Cdc42 family members Cdc42es2, Cdc42ep1, Cdc42ep2 or Cdc42ep4 was changed in the absence of Tgif1 during osteoblast adhesion and spreading (Fig. S3 A-D). These data indicate that the impaired spreading of Tgif1-deficient osteoblasts is unlikely caused by Cdc42 family members and that other factors are implicated in this process. We therefore quantified the expression of p21-activated-kinase (PAK) family members, who have been shown to play a role cell adhesion and migration as well as in actin nucleation in neurons (Liu et al., 2010) (Kreis and Barnier, 2009). While expression of PAK 1, 2 and 4 was unchanged in Tgif1-deficient osteoblasts during adhesion and spreading (Fig. S3 E-G), expression of PAK3 was strongly increased in Tgif1-deficient osteoblasts compared to control cells at the mRNA and protein level prior to and 60 min after adhesion (Fig. 3 A and B). Confirming this observation *in vivo*, expression of PAK3 mRNA showed a trend towards an increase in bones from *Tgif1^-/-^* mice compared to control littermates (Fig. 3C). These findings suggest that the impaired adhesion and spreading of Tgif1-deficient osteoblasts might be related to a deregulated abundance of PAK3.

**Figure 3.**
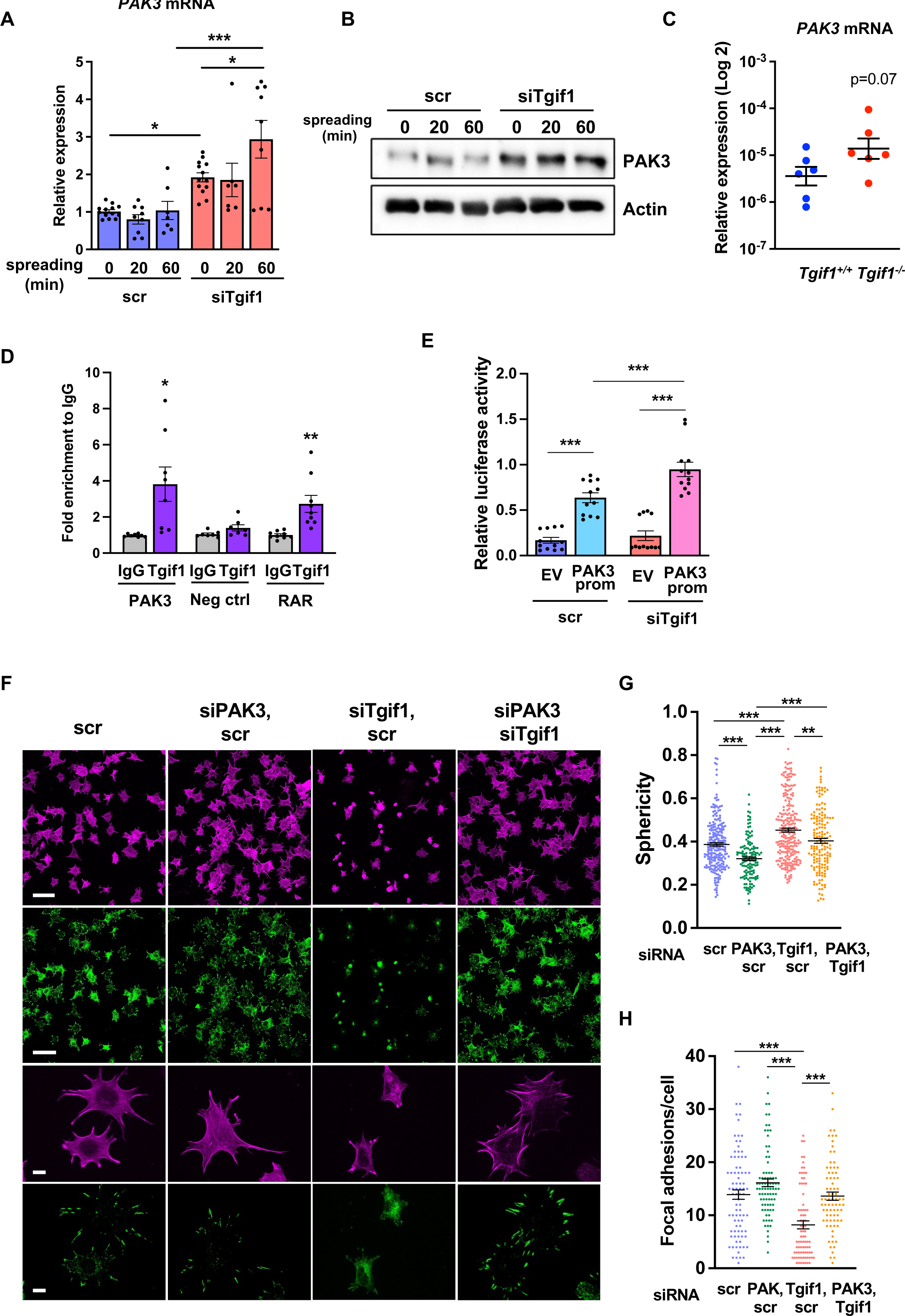
Silencing of PAK3 restores cell spreading and focal adhesion formation in Tgif1-deficient cells. **(A)** *PAK3* mRNA expression in OCY454 cells transfected with siRNA targeting Tgif1 (siTgif1) or scrambled control (scr) siRNA before (0) and after 20 and 60 minutes of spreading on Col-I coated slides. n=9, 1-way ANOVA with Tukey’s multiple comparison test, *p<0.05, ***p<0.001. **(B)** Representative images of immunoblots demonstrating the abundance of PAK3 expression upon silencing of Tgif1. Actin was used as a control. **(C)** *PAK3* mRNA expression in tibiae of 12-week-old *Tgif1^+/+^*(n=6) and *Tgif1^-/-^* (n=6) mice. **(D)** Tgif1 binding to the predicted site of the PAK3 promoter in OCY454 cells analyzed by ChIP and quantified as fold enrichment to the relative IgG control. Negative and positive (RAR) controls were used as indicated. n=8, unpaired t-test with Welch’s correction, *p<0.05, **p<0.01 vs. respective IgG control. **(E)** Tgif1-deficient (siTgif1) or control (scr) OCY454 cells were transfected with renilla plasmid and a pGL3 plasmid (EV) or a pGL3 plasmid containing a 2.3 kb fragment of the rat PAK3 promoter upstream of the luciferase gene. The promoter activity was quantified using luciferase assays and presented as relative luciferase activity (luciferase/renilla). 1-way ANOVA, Tukey’s multiple comparison test, ***p<0.001. **(F-H)** OCY454 cells were transfected alone or in combinations with siTgif1 for 48h, siPAK3 for 24 h and scrambled (scr) control. **(F)** Cells were allowed to adhere on Col-I coated slides for 60 minutes. Formation of focal adhesions was visualized by paxillin staining (green) and actin cytoskeleton by phalloidin staining (magenta). Scale bars indicate 100 µm (two upper rows) or 10 µm (two lower rows). **(G)** Quantification of cell sphericity using IMARIS. **(H)** Quantification of the number of mature focal adhesions per cell using the Image J software. n=4 independent experiments in which individual cells were analyzed. 1-way ANOVA, Tukey’s multiple comparison test, **p<0.01 ***p<0.001.

To further investigate the regulation of PAK3 expression by Tgif1, we performed an *in silico* analysis of the PAK3 promoter and identified 3 putative Tgif1 binding sites (Fig. S4A). Since only one site was species-conserved between rat and mouse, we verified the binding of Tgif1 to this site using chromatin immunoprecipitation. Indeed, Tgif1 bound to the predicted promoter binding site and to a site of the RAR promoter as positive control (Zhang et al., 2009), but not to a DNA sequence lacking predicted Tgif1 binding site as negative control (Fig. 3D), suggesting a regulation of the PAK3 promoter activity by Tgif1. To address this question, Tgif1 expression was silenced in OCY454 osteoblast-like cells using siRNA followed by transfection of the cells with a 2.3kb fragment of the rat PAK3 promoter upstream of the luciferase gene. Silencing of Tgif1 lead to a significant increase in luciferase activity compared to control (Fig. 3E) demonstrating that Tgif1 is a transcriptional repressor of the PAK3 promoter.

To determine if an elevated abundance of PAK3 due to Tgif1-deficiency impairs osteoblast spreading, PAK3 expression was silenced in Tgif1-deficient OCY454 cells using siRNA (Fig. S4B). Indeed, attenuating PAK3 expression restored the Tgif1 deficiency-mediated impaired cell spreading (Fig. 3F). Since cells are less spherical upon spreading, PAK3 silencing reduced the Tgif1 deficiency-dependent gain in cell sphericity and restored the number of focal adhesions per cell that were reduced by Tgif1 deficiency to the level of control (Fig. 3G and H). Together, these findings demonstrate that in the absence of Tgif1, the PAK3 promoter is de-repressed, leading to an increased transcriptional activity and PAK3 expression. A higher abundance of PAK3 in turn suppresses focal adhesion formation and osteoblast spreading.

### Tgif1 expression increases during osteoblast spreading via activation of ERK1/2 and AP1 signaling pathways

To further investigate the role of Tgif1 during osteoblast spreading, we quantified Tgif1 expression during this process. Both, Tgif1 mRNA and protein expression were significantly increased within 20 minutes of spreading with a further increase after 60 minutes in calvarial osteoblasts (Fig. 4A and B) and in OCY454 cells (Fig. 4C and D). To determine if the increase in Tgif1 abundance is transcriptionally regulated, osteoblasts were treated with the transcription inhibitor Actinomycin-D during spreading, which fully prevented the increase in Tgif1 mRNA expression (Fig. 4E).

**Figure 4.**
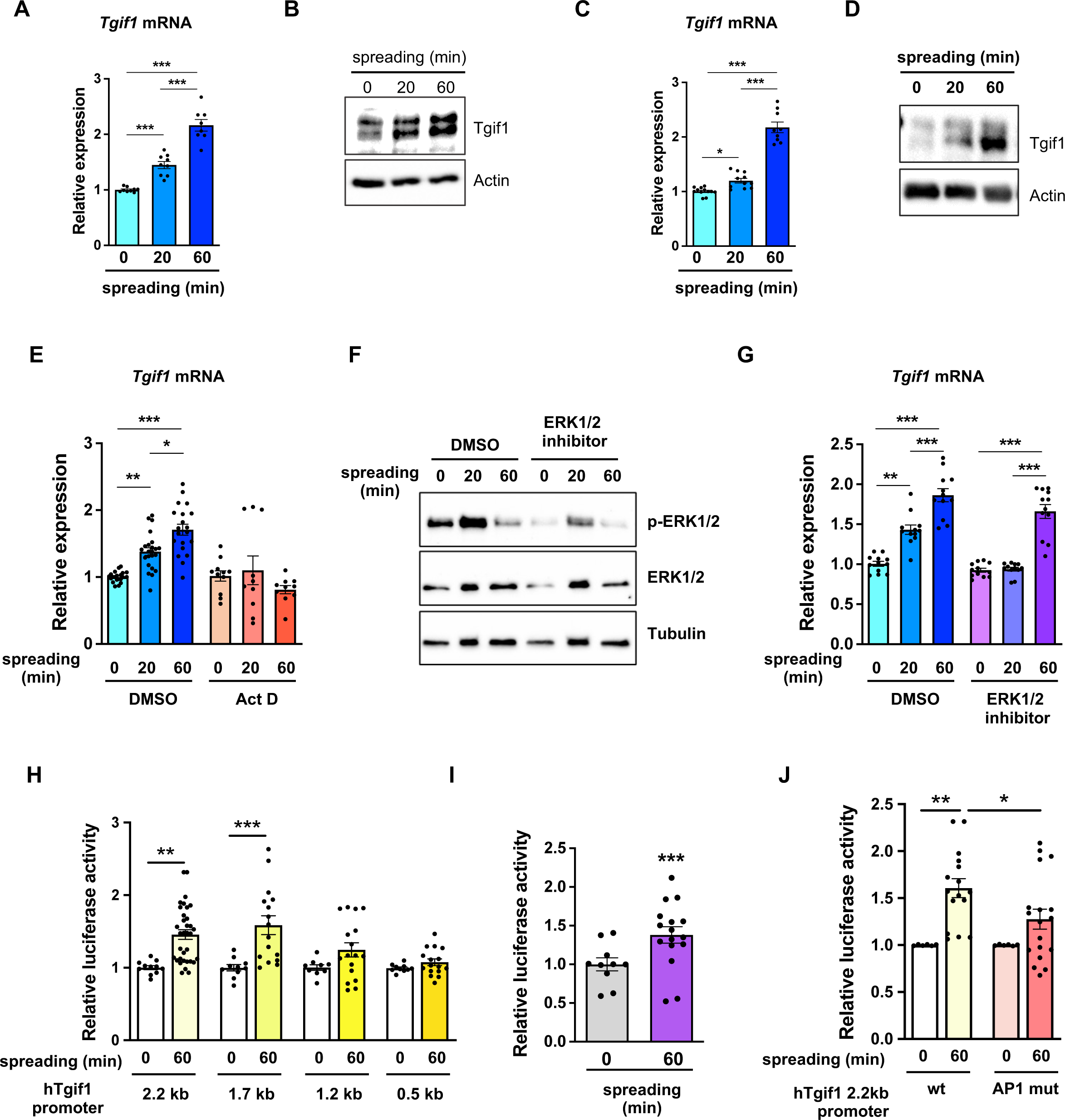
Tgif1 expression is increased during cell spreading via ERK and AP1 signaling pathways. **(A)** Quantification of *Tgif1* mRNA expression and **(B)** representative images of immunoblots of Tgif1 expression in calvarial osteoblasts before (0 min) and after 20 and 60 minutes of spreading. Actin was used as control. **(C)** *Tgif1* mRNA expression and **(D)** representative images of immunoblots of Tgif1 expression in OCY454 cells before (0 min) and after 20 and 60 minutes spreading on Col-I coated slides. Actin was used as control. **(E)** OCY454 cells were treated with DMSO as control or with 5 µM Actinomycin D for 15 minutes prior to adherence on Col-I coated slides for 20 and 60 minutes. *Tgif1* mRNA was quantified before (0 min) and during spreading. **(F)** Representative immunoblot images demonstrating the efficiency of ERK1/2 inhibitor SCH772984 (1µg/ml) to prevent ERK1/2 phosphorylation (n=4). **(G)** Quantification of *Tgif1* mRNA expression during spreading after 15 minutes pre-treatment with DMSO (control) or with the ERK inhibitor SCH772984. **(H)** Quantification of the Tgif1 promoter activity during cell spreading. OCY454 cells were transfected with a luciferase reporter plasmid encoding a 2.2 kb fragment of the Tgif1 promoter or progressive truncations thereof (1.7, 1.2 and 0.5 kb) along with a plasmid encoding renilla firefly as control. Upon spreading for 60 minutes, promoter activity was quantified using a dual luciferase reporter gene assay and presented as normalized luciferase activity (luciferase/renilla). **(I)** Quantification of the AP1 transcriptional activity during cell spreading. OCY454 cells were transfected with a 6X-TRE-luciferase reporter to determine AP-1 activity along with a plasmid encoding renilla firefly as control. Upon spreading for 60 minutes, promoter activity was quantified using a dual luciferase reporter gene assay and presented as normalized luciferase activity (luciferase/renilla). **(J)** Quantification of the activity of the wild-type (wt) Tgif1 2.2 kb promoter and of the same fragment bearing a mutant AP1 binding site (AP1 mut) during cell spreading. OCY454 cells were transfected either with a luciferase reporter plasmid encoding a wild-type (wt) 2.2 kb fragment of the Tgif1 promoter or with the same promoter in which the AP1 binding site has been mutated (mut) along with a plasmid encoding renilla firefly as control. Upon spreading for 60 minutes, the promoter activity was quantified using a dual luciferase reporter gene assay and presented as normalized luciferase activity (luciferase/renilla). 1-way ANOVA, Tukey’s multiple comparisons test, *p<0.05, **p<0.01, ***p<0.001 vs. respective control.

Since binding of cells to collagen I-coated surfaces activates the ERK1/2 pathway (Emerson et al., 2009; Fincham et al., 2000), we investigated the potential implication of ERK1/2 signaling in the increase of Tgif1 gene expression during osteoblast spreading. Experimentally, osteoblasts were treated with an ERK1/2 inhibitor, which greatly attenuated ERK1/2 phosphorylation after 20 min and 60 min of osteoblast spreading (Fig. 4F). Although ERK1/2 inhibition prevented the increase of Tgif1 mRNA expression during the first 20 min of osteoblast spreading, it failed to suppress Tgif1 mRNA expression 60 min after initiation of osteoblast spreading (Fig. 4G). This finding suggests that the increase in Tgif1 mRNA expression at early stages of osteoblast spreading is mediated by ERK1/2 signaling while it is independent of ERK1/2 signaling at later stages of osteoblast spreading.

To elucidate the molecular mechanisms underlying the activation of Tgif1 transcription at later stages of osteoblast spreading, we performed progressive truncations of the 2.2 kb fragment of the human Tgif1 promoter (Saito et al., 2019). While the 2.2 kb and 1.7 kb fragments of the human Tgif1 promoter were fully activated after 60 minutes of osteoblast spreading, no activation of the 1.2 kb and 0.5 kb fragments was observed (Fig. 4H), suggesting that the regulatory promoter region that is activated during osteoblast spreading must be located within the 500 bp between the 1.7 kb and the 1.2 kb promoter fragments.

Recently we reported the presence of an AP1 binding site in this region of the Tgif1 promoter (Saito et al., 2019), suggesting that AP1 signaling might be implicated in this process. Since the role of AP1 signaling in the context of cell spreading has not been fully elucidated, we first determined the activation of an AP1-responsive promoter element during osteoblast spreading. The findings demonstrate that AP1-signaling is activated 60 minutes after initiation of osteoblast spreading (Fig. 4I), indicating a potential implication of AP1 signaling during advanced stages of osteoblast spreading. To further test this hypothesis, osteoblasts were transfected either with a plasmid encoding the wild-type Tgif1 promoter bearing an AP1 binding site or with a plasmid in which the AP1 binding site was disabled by site-specific mutation (Saito et al., 2019). Although the wild-type promoter was fully activated after 60 min of osteoblast spreading, the Tgif1 promoter bearing the disabled AP1 binding site was not activated (Fig. 4J). These findings suggest that AP1 signaling is involved in the activation of Tgif1 gene expression during later stages of osteoblast spreading.

### Tgif1 deficiency leads to fewer and less active osteoblasts during bone regeneration

Bone has a regenerative capacity to heal upon fracture. In fulfilling this function, osteoblast adherence and spreading are crucial features since osteoblasts migrate to sites of bone repair for subsequent matrix production and regeneration (Thiel et al., 2018; Dirckx et al., 2013). Fracture healing involves various cell types and cellular processes (Einhorn and Gerstenfeld, 2015). One important component is the activation of periosteal cells to migrate alongside vessels that sprout into the fracture zone (Maes et al., 2010). Periosteal cells become active osteoblasts that produce a cartilage template, which then ossifies and forms new trabeculae to consolidate the fracture gap (Duchamp De Lageneste et al., 2018). Since this process involves adherence and spreading of osteoblasts, we hypothesized that Tgif1 might be important in the context of bone repair. To test this hypothesis, we performed open mid-shaft tibia fractures in 10-weeks old adult male mice bearing a germline deletion of Tgif1 and control littermates, followed by fracture stabilization using an intramedullary pin. Three weeks after fracture, newly formed trabeculae were densely occupied by cuboidal matrix-producing osteoblasts in control animals (Fig. 5 A-E). In contrast, in mice bearing a germline deletion of Tgif1, the callus was less ossified and the trabeculae in the repair zone were sparsely occupied by fewer, thinner, and less-active osteoblasts (Fig. 5 A-F). These findings demonstrate that Tgif1 is important for bone regeneration.

**Figure 5:**
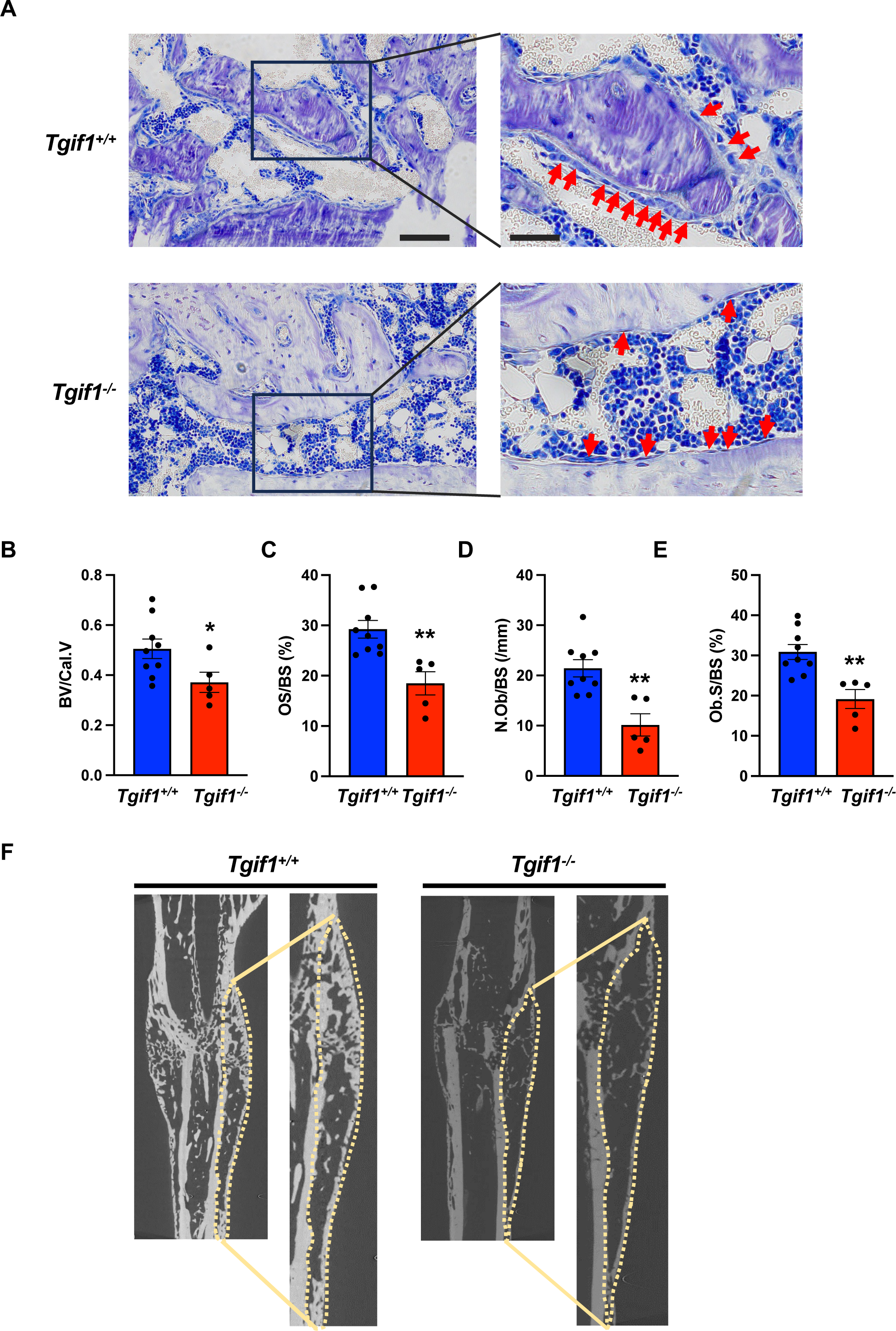
Tgif1 deficiency causes a reduced number and activity of osteoblasts during bone repair. **(A)** Representative images of Toluidine blue-stained sections of bones from male *Tgif1^+/+^*(n=9) and *Tgif1^-/-^* mice (n=5) that received an open fracture of the midshaft tibia at 10 weeks of age followed by bone healing for 3 weeks upon intramedullary stabilization. Arrows indicate osteoblasts. Scale bars indicate 100 µm (left panels) and 50 µm (right panels). **(B - E)** Histological sections were used to quantify the histomorphometric parameters **(B)** bone volume per callus volume (BV/Cal.V), **(C)** osteoid surface per bone surface (OS/BS), **(D)** number of osteoblasts per bone surface (N.Ob/BS) and **(E)** osteoblast surface per bone surface (Ob.S/BS). **(F)** Representative µCT images of the tibiae of *Tgif1^+/+^* and *Tgif1^-/-^* mice 21 days after fracture. Unpaired t-test, *p<0.05, **p<0.01 vs. *Tgif1^+/+^*.

### Activation of bone surfaces by PTH is attenuated in the absence of Tgif1 in osteoblasts

Intermittent administration of PTH augments bone formation, leading to a remodeling-based increase in bone mass, bone mineral density and a consecutive decrease in fracture rate in humans (Neer et al., 2001; Taipaleenmäki et al., 2022; Baron and Hesse, 2012). This anabolic function requires multiple alterations at the cellular level. For instance, PTH induces the differentiation of mesenchymal precursor cells and reverts quiescent bone lining cells into active osteoblasts (Nishida et al., 1994; Dobnig and Turner, 1995; Kim et al., 2012). Furthermore, PTH increases the number and bone-forming capacity of mature osteoblasts (Mizoguchi and Ono, 2021; Jilka et al., 2009).

Recently, we identified Tgif1 as a PTH target gene in osteoblasts that is necessary for the full PTH-mediated bone mass accrual (Saito et al., 2019). In addition, the site of the Tgif1 promoter known to be activated by PTH signaling (Saito et al., 2019), is identical with the site identified here that is activated by AP1 signaling during osteoblast adhesion (Fig. 4J). In this study we also uncovered that Tgif1 is a physiological regulator of osteoblast adherence, spreading and migration. Collectively, this evidence let us to propose that Tgif1 might also be implicated in the PTH-mediated bone surface activation.

To determine whether Tgif1 is implicated in the PTH-mediated activation of bone surfaces by osteoblasts, we performed histomorphometric analysis of bone sections from mice bearing a germline deletion of Tgif1 and control littermates that were injected with PTH or vehicle control. Consistent with the findings reported by others and in support of our hypothesis, PTH treatment greatly increased the number of active osteoblasts and induces the acquisition of a cuboidal shape and consequently the percentage of active bone surfaces (Kousteni and Bilezikian, 2008; Tam et al., 1982), which all occurred at a much lesser extent in mice lacking Tgif1 (Fig. 6A and B). To determine whether this effect is osteoblast-autonomous, we performed histomorphometric analysis of bones obtained from mice in which Tgif1 deletion was targeted to mature osteoblasts using the Dmp1-promoter (Bivi et al., 2012). Indeed, while PTH treatment induced osteoblast recruitment, adaptation of a cuboidal morphology of matrix-producing osteoblasts and a profound activation of bone surfaces by osteoblasts in control animals, these effects were much less pronounced in mice lacking Tgif1 in osteoblasts (Fig. 6C and D). These findings therefore demonstrate that PTH-dependent bone surface activation is attenuated in the absence of Tgif1 in osteoblasts.

**Figure 6.**
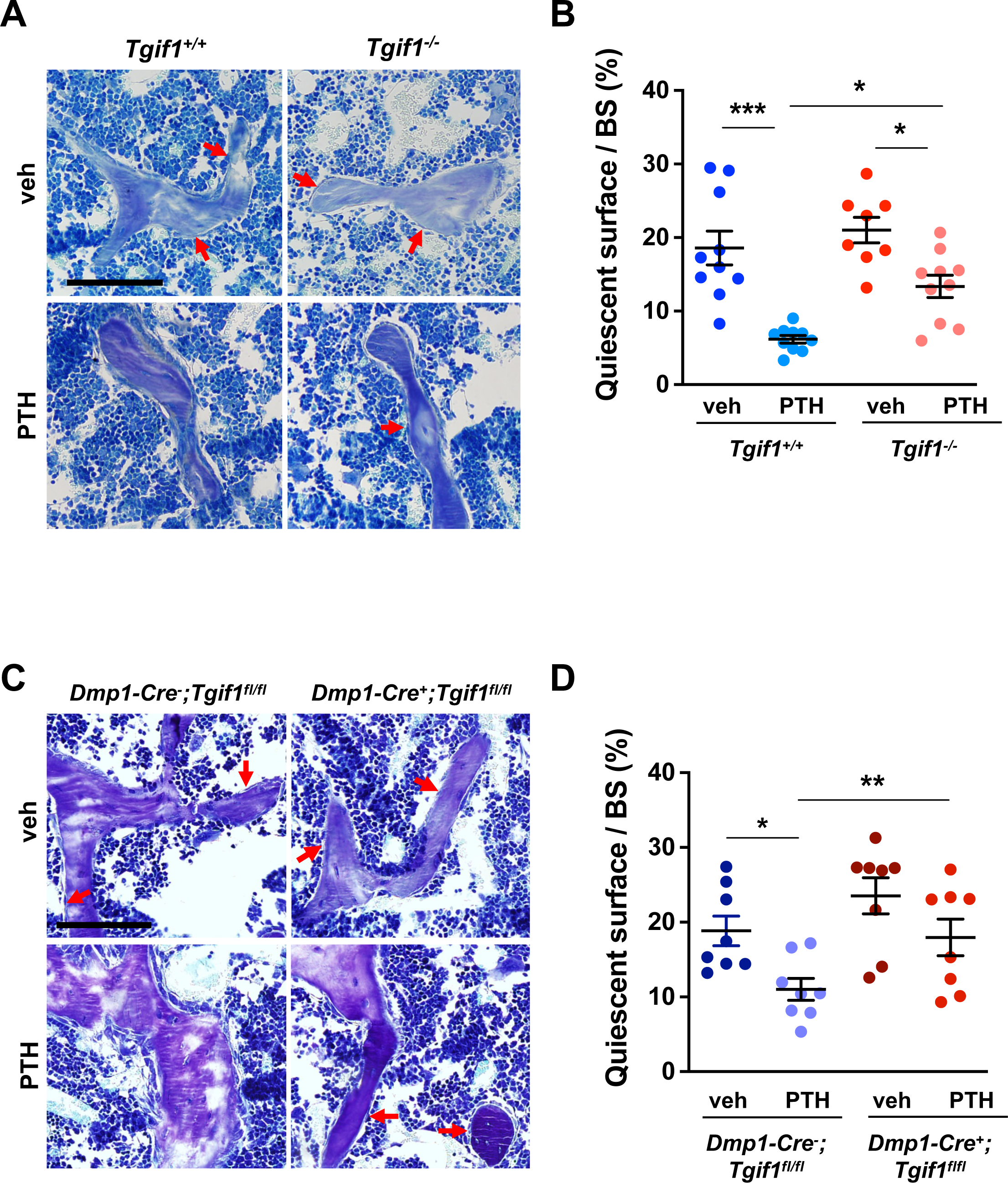
PTH is impaired in activating quiescent bone surfaces in *Tgif1*-deficient mice. **(A)** Toluidine blue staining of tibiae from 12-week-old *Tgif1^+/+^* and *Tgif1^-/-^* mice treated with vehicle (veh) or PTH. Quiescent surfaces are indicated by red arrows. Representative images are shown. Scale bar indicates 100 µm. **(B)** Quantification of the percentage of quiescent surfaces per bone surface (BS). *Tgif1^+/+^*, veh n=10; *Tgif1^+/+^*, PTH n=10, *Tgif1^-/-^*, veh n=8; *Tgif1^-/-^*, PTH n=10. (**C**) Toluidine blue staining of tibiae from 12-week-old *Dmp1-Cre^-^;Tgif1^fl/fl^*and *Dmp1-Cre^+^;Tgif1^fl/fl^* mice treated with vehicle (veh) or PTH. Quiescent surfaces are indicated by red arrows. Representative images are shown. Scale bar indicates 100 µm. **(D)** Quantification of the percentage of quiescent surfaces per bone surface (BS). *Dmp1-Cre^-^;Tgif1^fl/fl^*, veh n=8; *Dmp1-Cre^-^;Tgif1^fl/fl^*, PTH n=8, *Dmp1-Cre^+^;Tgif1^fl/fl^*, veh n=8; *Dmp1-Cre^+^;Tgif1^fl/fl^*, PTH n=8. 1-way ANOVA with Tukey’s multiple comparison test, *p<0.05, **p<0.01.

### PTH facilitates spreading of osteoblasts via Tgif1-PAK3 signaling

The observation that Tgif1 is crucial for osteoblast adherence, spreading and migration *in vitro* as well as for the PTH-mediated activation of bone surfaces suggests that PTH may facilitate the spreading of osteoblasts and therefore the decrease in cell sphericity via Tgif1. To test this hypothesis, OCY454 cells were treated with PTH or vehicle control during spreading. Immunofluorescence staining and quantification of cell sphericity revealed that PTH supports osteoblast spreading (Fig. 7A and B). Furthermore, PTH treatment of cells in which Tgif1 expression was silenced failed to induce the spreading of osteoblasts (Fig. 7A and B), demonstrating that the PTH-mediated osteoblast spreading indeed depends on Tgif1.

**Figure 7.**
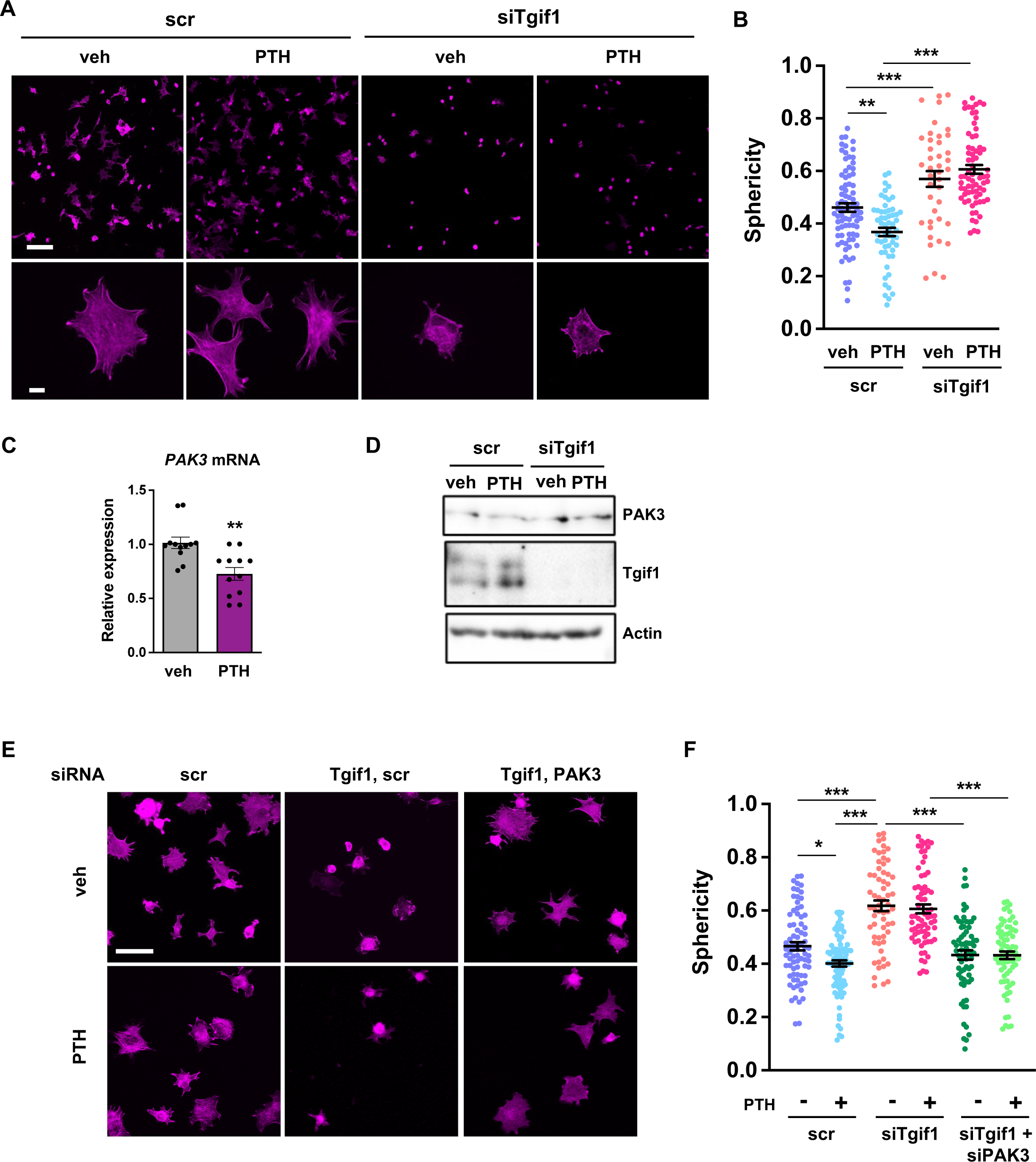
PTH promotes cell spreading via Tgif1-PAK3 signaling. **(A)** OCY454 cells were transfected with siRNA against Tgif1 (siTgif1) or scramble control siRNA (scr) and treated with vehicle (veh) or PTH for 4h. Cells were allowed to adhere on Col-I coated slides for 60 minutes. The cytoskeleton was visualized by phalloidin staining (magenta). Scale bars indicate 100 µm (upper panel) or 10 µm (lower panel). **(B)** Cell sphericity was quantified using IMARIS software. 1-way ANOVA with Tukey’s multiple comparisons test was applied, **p<0.01, ***p<0.001. **(C)** *PAK3* mRNA expression after 4 hours of PTH treatment. Unpaired t-test, **p<0.01 vs. veh. **(D)** Representative image of immunoblots demonstrating PAK3 and Tgif1 protein abundance upon PTH treatment in cells transfected with siTgif1 or scr. Actin was used as a loading control. **(E)** OCY454 cells were transfected with scr or Tgif1 siRNA for 48h and PAK3 or scr siRNA for 24 h and treated with vehicle (veh) or PTH for 4h. Cells adhered on Col-I coated slides for 60 minutes. The cytoskeleton was visualized by phalloidin staining (magenta). Scale bar indicates 50 µm. **(F)** Cell sphericity was quantified using IMARIS software. n=4, 1-way ANOVA, Tukey’s multiple comparison test, *p<0.05, ***p<0.001.

Since PAK3 expression is increased in the absence of Tgif1 and impairs focal adhesion formation and osteoblast spreading, we investigated the possibility that PTH decreases PAK3 gene expression via Tgif1. Indeed, PAK3 mRNA expression was reduced upon PTH treatment (Fig. 7C). In addition, PTH treatment increased Tgif1 expression and reduced PAK3 protein abundance in control cells but not in osteoblasts in which Tgif1 expression was silenced by siRNA (Fig. 7D). Next, we examined osteoblast spreading upon PTH treatment in the absence of Tgif1 alone or in combination with loss of PAK3. Immunofluorescence staining and quantification of cell sphericity revealed that silencing Tgif1 expression impaired osteoblast spreading induced by PTH treatment, which was restored by the concomitant silencing of PAK3 expression (Fig. 7E and F). Furthermore, lack of both, Tgif1 and PAK3 prevented PTH-induced decrease in cell sphericity. These findings indicate that an increased abundance of PAK3 in response to Tgif1-deficiency impairs PTH-induced spreading of osteoblasts and that PTH facilitates osteoblast spreading via Tgif1-PAK3 signaling.

In summary, our results reveal that Tgif1 suppresses PAK3 expression in osteoblasts via the ERK1/2 and AP1 signaling pathways, which is important for osteoblasts to form focal adhesions, to spread on bone surfaces, and to migrate. This mechanism is implicated in bone regeneration and in the pharmacological effects of PTH treatment and is therefore of translational relevance.

## Discussion

This study revealed that Tgif1-deficient osteoblasts exhibit an altered morphology, reduced adherence to collagen type I-coated surfaces, impaired migration capacity, and decreased spreading compared to control cells. These defects in Tgif1-deficient osteoblasts are associated with compromised focal adhesion formation and increased expression of PAK3. The findings further demonstrate that Tgif1 regulates PAK3 expression through transcriptional repression and that elevated PAK3 abundance contributes to the impaired osteoblast spreading observed in Tgif1-deficient cells. Additionally, Tgif1 is shown to participate in the activation of bone surfaces in the context of bone regeneration and PTH treatment since in the absence of Tgif1, bone regeneration- and PTH-induced activation of bone surfaces is attenuated. Mechanistically, PTH promotes osteoblast spreading via Tgif1-PAK3 signaling, with PAK3 expression being reduced by PTH treatment in a Tgif1-dependent manner. Overall, this study emphasizes the importance of Tgif1 in regulating osteoblast morphology, adherence, and migration through the modulation of PAK3 expression, providing novel and translationally relevant mechanistic insights into the regulation of the cytoskeletal architecture of osteoblasts.

Both, bone repair and PTH treatment are frequent and clinically relevant events. Fractures often occur in the context of high energy accidents, however, bone fragility syndromes such as osteoporosis also cause fractures upon minimal or no trauma (Malluche et al., 2013; Löfman et al., 2009). Bone regeneration, a complex process involving osteoblast recruitment, matrix production, and tissue regeneration, is compromised in Tgif1-deficient mice. Activation of bone surfaces by osteoblasts, a critically important aspect of bone repair, is significantly reduced in the absence of Tgif1, leading to a reduced callus ossification. In addition, Tgif1-deficient mice also have a reduced number and activity of osteoclasts mice (Saito et al., 2019) and therefore an attenuated bone remodeling which might also contribute to an impaired bone healing. Nevertheless, the results presented here indicate that Tgif1 is important for osteoblast recruitment to the site of repair and subsequent bone surface activation, which extends the established function of Tgif1 beyond the regulation of osteoblast differentiation and activity.

Albeit mechanistically distinct, bone repair and PTH treatment have in common that osteoblasts are an important component for the successful execution of bone regeneration and the PTH pharmacological effects. In particular, the guided spatial relocation of activated osteoblasts is a critical feature to exert the respective functions at the right sites in a three-dimensional context since formation of new bone, regardless of whether in the cortical- or trabecular compartment, occurs in a timely but also in a spatially controlled manner to yield structurally supportive newly formed bone tissue. Various types of cells and tissues of the local microenvironment produce a plethora of signaling factors, which are induced by fractures or PTH treatment. However, little is known about signaling pathways in osteoblasts that receive and integrate these diverse external cues to facilitate the adherence, spreading and movement of osteoblasts in a coordinated manner. In this context, this study identified the functional importance of PAK3 in Tgif1-deficient osteoblasts. PAK3 expression was increased in the absence of Tgif1, leading to impaired focal adhesion formation and reduced osteoblast spreading. Silencing PAK3 expression restored osteoblast spreading in Tgif1-deficient cells, demonstrating that PAK3 plays a critical role in mediating the effects of Tgif1 on osteoblast morphology. This work therefore contributes to the better understanding of osteoblast dynamics by identifying Tgif1 and PAK3 as downstream signaling regulators that are important for osteoblasts to fulfill motility. Furthermore, the work underscores the physiological relevance of spatial osteoblast dynamics as integral component of bone remodeling and maintenance of skeletal integrity.

Tgif1 recently emerged as a key determinant of osteoblast activity, bone mass maintenance and essential component of the bone anabolic PTH signaling pathway (Haider et al., 2020; Saito et al., 2019) This study also focused on investigating the role of PTH in regulating osteoblast spreading and revealed that PTH treatment facilitated osteoblast spreading via Tgif1-PAK3 signaling. PTH treatment reduced PAK3 expression and increased Tgif1 expression, leading to enhanced osteoblast spreading. Silencing Tgif1 expression impaired PTH-induced osteoblast spreading, which was restored by simultaneously silencing PAK3 expression. These new findings on PTH-Tgif1-PAK3 signaling provide novel insights into the function of PTH treatment of bone remodeling and the implication of Tgif1 in this process. As part of being required for bone mass accrual in response to PTH treatment (Saito et al., 2019), Tgif1 also contributes to the PTH-induced osteoblast recruitment and subsequent movement to sites where new bone needs to be formed. This adds a spatial component to a functional feature and demonstrates the complexity of bone remodeling and PTH anabolic actions.

This work has the limitation that the findings on the role of Tgif1 in osteoblast spreading and migration were obtained by *in vitro* assays, which do not necessarily fully reflect *in vivo* situations. Furthermore, limited evidence exists that the role of Tgif1-PAK3 signaling uncovered *in vitro* also plays a crucial role *in vivo*. Thus, future *in vivo* studies need to further validate our findings for instance by determining PAK3 expression in *Tgif1^-/-^* mice in fractures and upon PTH treatment as well as by pharmacologically suppressing PAK3 expression in Tgif1-deficient mice.

In summary, these findings highlight the pivotal role of Tgif1 in regulating osteoblast morphology, adherence, and migration. Notwithstanding the possibility that other mechanisms exist, our data reveal the involvement of PAK3 in mediating the effects of Tgif1 on osteoblast spreading. The dysregulation of these processes due to Tgif1 deficiency has profound implications for bone remodeling, bone regeneration, and the pharmacological effects of PTH. Reaching a better understanding of the intricate interplay between PTH, Tgif1 and PAK3, provides valuable insights into the molecular mechanisms governing osteoblast function and could have translational relevance in bone-related disorders and therapies targeting bone health.

## Materials and methods

### Animal models

Mice with germ-line deletion of Tgif1 or loxP-flanked Tgif1 loci have been reported previously (Shen and Walsh, 2005). To delete Tgif1 in mature osteoblasts and osteocytes, mice expressing the Cre recombinase under the control of the 8kb fragment of the murine Dentin matrix protein 1 (Dmp1-Cre^Tg^) (Bivi et al., 2012) promoter were crossed with mice in which exons 2 and 3 of the Tgif1 gene were flanked by loxP sites (Tgif1^fl/+^). Since no bone phenotype was observed in *Dmp1-Cre^+^*; *Tgif1^+/+^* mice, *Dmp1-Cre^-^; Tgif1^fl/fl^* mice were used as controls (Saito et al., 2019). In anabolic studies, parathyroid hormone (PTH 1-34; 100 µg/kg of body weight, Biochem) was administered in 8-week-old male mice intraperitoneally 5 times a week for 3 weeks. To induce conditions of bone healing, 10-week-old male mice of the genotypes *Tgif1^+/+^* or *Tgif1^-/-^* were subject to an open fracture of the tibial midshaft, which was stabilized by an intramedullary pin (Schindeler et al., 2008). Three weeks later, mice were sacrificed with subsequent histological examination of the bone tissue. All animal experiments were approved by the local authorities.

### Histology and histomorphometry

Mouse tibiae were collected and fixed in 3.7% PBS-buffered formaldehyde. For histomorphometric analysis, tibiae were embedded in methylmethacrylate. Toluidine blue staining was performed on 4 µm sagittal sections. Quantitative bone histomorphometric measurements were performed on von Kossa and Toluidine blue stained sections using the OsteoMeasure system (OsteoMetrics). For the quantification of quiescent surfaces (Qs) only surfaces covered by flat cells of intense blue staining without osteoid deposition were considered.

### Micro-computed tomography

Three-dimensional (3D) visualization of the callus formed at the fracture site was performed using micro-computed tomography (μCT). By day 21 upon open fracture of the tibial midshaft, mice were sacrificed and whole legs were harvested followed by fixation in 3,7 % paraformaldehyde. Intramedullary pins were removed prior to imaging to avoid artifacts. Fractured tibiae were scanned using high-resolution μCT with a fixed isotropic voxel size of 10 μm (70 kV at 114 μA, 400 ms integration time; Viva80 micro-CT; Scanco Medical AG). To visualize bone tissue, the threshold value was determined at 326 mg/cm^3^ hydroxyapatite based on Hounsfield units and a phantom with a linear hydroxyapatite gradient (79–729 mg/cm^3^). To determine the abundance of mineralized bone within the middle of the callus area, a region of interest was defined and magnified for better visualization.

### Isolation of primary osteoblasts and cell culture

To obtain calvarial osteoblasts, mouse calvariae were dissected from 1–3 days old neonatal mice and digested sequentially 5 times for 25 minutes in α-MEM (Gibco) containing 0.1% collagenase (Roche) and 0.2% dispase (Roche). Cells obtained from fractions 2–5 were combined according to the genotype and expanded in α-MEM containing 10% FBS (Gibco), 100 U/ml penicillin, and 100 µg/ml streptomycin (P/S, Thermo Fisher Scientific). Long bone osteoblasts were isolated from femora and tibiae of 8-week-old mice. After removing muscles in sterile PBS, bone marrow was flushed, and bones were cut in small pieces. Bone pieces were digested with 0.1% collagenase for 2 hours at 37 °C and plated in α-MEM containing 10% FBS and P/S. The OCY454 cell line (RRID:CVCL_UW31) was kindly provided by Dr. Paola Divieti Pajevic (Department of Molecular and Cell Biology, Boston University, Boston, USA) and cultured in α-MEM with 10% FBS, 100 U/ml penicillin, and 100 µg/ml streptomycin at 33°C to permit proliferation and maintain osteoblast phenotype, and at 37°C for the experiments.

### 3D cell culture

Long bone osteoblasts were isolated as described above. Bone pieces were digested with 0.1% collagenase for 2 hours at 37 °C and plated in α-MEM containing 10% FBS and P/S. Once confluent, outgrowing osteoblasts were seeded on Cellmatrix Type 1A (Nitta Geratin) placed onto a Millicell® membrane (Sigma-Aldrich) in α-MEM with 10% FBS, 100 U/ml penicillin, and 100 µg/ml streptomycin, as reported previously (Uchihashi et al., 2013). After 24 hours, the membrane/gel/cells complexes were fixed in 1,5% formaldehyde, embedded in OCT and frozen. 10 µm sections were cut and stained with Hematoxylin/Eosin and for immunocytochemistry with phalloidin and DAPI as described below. Cell area and perimeter were analyzed from phalloidin-stained sections using the Fiji software.

### Cell spreading and migration assays

For adhesion and spreading assays, cells were incubated in α-MEM with 1% FBS for 4h and removed from the culture flasks by gentle trypsinization (Trypsin-EDTA 0.05%, Thermo Fisher Scientific) (Dejaeger et al., 2017). Cells were left to adhere for 20, 40, 60, 120 or 240 minutes on Collagen I-coated 8 well-slides (5*10^3^ cells/well) or plates (5*10^5^ cells/well). For qPCR, control cells were left in suspension. All spreading assays were performed in serum-free conditions at 37°C in duplicates or triplicates, which were combined prior to RNA or protein analysis. For the quantification of adhered cells, cells were fixed and stained with hematoxylin or incubated with Calcein–AM 2 μM (Thermo Fisher Scientific) for 60 minutes. For migration assays, cells were plated at a density of 7500 cells /cm^2^ on Collagen I-coated 6-well plates in culture medium (Dang and Gautreau, 2018). Cells adhered for 2-3 hours before the medium was changed to α-MEM containing 0.1% FBS. Live cell migration assay was captured using the Improvision LiveCell Spinning Disk microscope. For live cell imaging, 5 regions of interest/well were selected at time point 0 and images were captured every 10 minutes for 16 hours. Analysis of migration track length, velocity and meandering index was performed using the software Volocity 6.

### Cell treatment

For the inhibition of gene transcription, cells were incubated for 15 minutes with 5 µM of Actinomycin-D (Sigma). For the inhibition of ERK1/2 signaling, cells were incubated for 20 minutes with 1 µg/ml SCH772984 (Selleckchem) (Morris et al., 2013). For the *in vitro* PTH treatment, cells were incubated with 100 nM PTH (1-34, Biochem) in α-MEM containing 1% FBS for 4 hours prior to the spreading assays.

### RNA isolation and gene expression analysis

Total RNA was isolated using the RNeasy plus kit (Invitrogen) according to manufacturer’s instruction. cDNA was synthesized from total RNA using the ProtoScript First Strand cDNA Synthesis Kit (NEBioLabs). Quantitative real-time PCR was performed using SYBR Green (BioRad). The fold induction of each target gene was calculated using the comparative CT method (2^-ΔΔCt^). The C_t_ of each gene of interest was first normalized to the housekeeping gene GAPDH (ΔC_t_), and then to the control of the experiment (vehicle or DMSO treatment) to calculate the ΔΔC_t.,_ The oligonucleotide sequences used were the following:

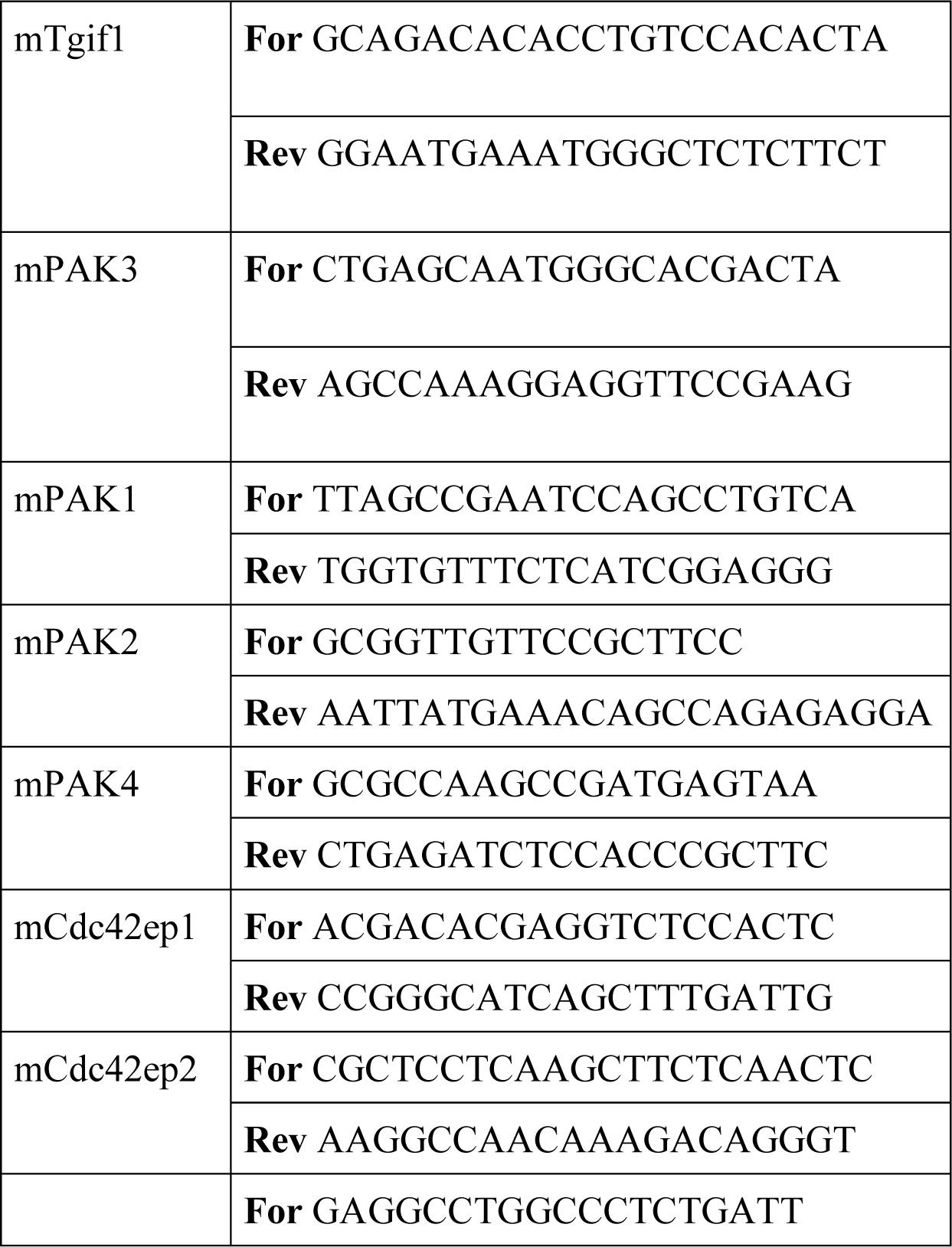
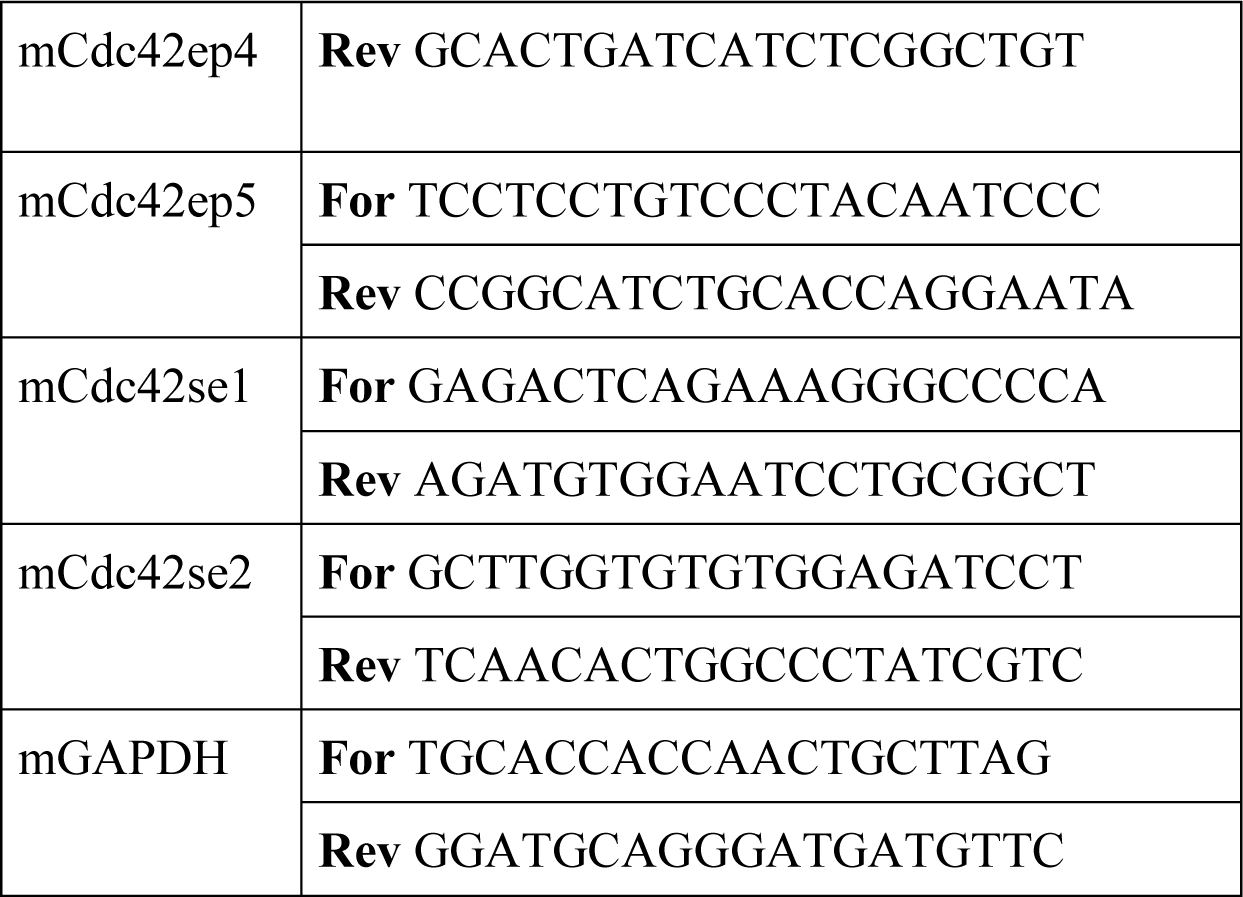

### Immunocytochemistry

Upon adherence of calvarial osteoblasts or OCY454 cells on Collagen I (Roche)-coated surfaces of 8-well chambers, cells were fixed in 1.5% formaldehyde and immune-stained with rabbit anti-Paxillin antibody (1:150, Abcam) or mouse anti-Talin antibody (1:100, Sigma), followed by an incubation with a 488-Alexa Fluor conjugated secondary antibody (1:500, Thermo Fisher Scientific). Hoechst 33258 (Thermo Fisher Scientific) was used at a 1:10000 dilution for nuclear staining. 633-Alexa Fluor conjugated phalloidin (Thermo Fisher Scientific) was used at a 1:20 dilution to stain actin. Cover slips were mounted with Fluoromount G (Southern Biotech).

### Image acquisition and quantification

Images were acquired using the Zeiss Apotome or the confocal Leica SP5 and SP8 microscopes with 20X or 63X objectives. Images for the analysis of focal adhesions were quantified with the Image J software as described previously (Horzum et al., 2014). Images for 3D quantification were elaborated by IMARIS software (BITPLANE, Oxford). The Z-stacks obtained by confocal microscopy were rebuilt by IMARIS and the parameters of interest were quantified after removal of the background noise. The value of the sphericity for each cell was provided automatically by the software after cell selection. Quantification of images was performed using raw data. Representative images shown have been optimized equally and only for brightness and contrast.

### Immunoblot analysis

Samples were lysed in low salt RIPA buffer pH 7.5 (50 mM Tris base, 150 mM NaCl, 0.5% Nonidet P-40, 0.25 % Sodium deoxycholate,) supplemented with complete protease and phosphatase inhibitors (Roche). Lysates were denaturated in Laemmli Sample Buffer (BioRad), resolved by SDS-PAGE and transferred onto nitrocellulose membranes (GE Healthcare). Membranes were incubated with the following primary antibodies: rabbit anti-Tgif1 antibody (Abcam, 1:500), rabbit anti-PAK3 antibody (Cell Signalling, 1:300), rabbit anti-phospho ERK1/2 antibody (Cell Signalling, 1:1000), mouse anti-ERK1/2 antibody (Cell Signalling, 1:1000), mouse anti-tubulin antibody (Sigma, 1:2000), mouse anti-actin antibody (Millipore, 1:5000), rabbit anti-integrin β1 (Cell Signaling, 1:500), rabbit-anti FAK (Cell Signaling, 1:500), rabbit-anti p-FAK (Cell Signaling, 1:500), rabbit-anti paxillin (Cell Signaling, 1:500), rabbit-anti p-S83 paxillin (Cell Signaling, 1:500), mouse anti-src (Cell Signaling, 1:500), rabbit anti-phospho src (Cell Signaling, 1:500), rabbit anti-p38 (Cell Signaling, 1:500), rabbit anti-phospho p38 (Cell Signaling, 1:500), rabbit-anti LRG5 (Cell Signaling, 1:500). Membranes were washed with TBST and incubated either with anti-mouse or anti-rabbit IgG horseradish peroxidase-conjugated secondary antibody (Promega, 1:10000). Chemiluminescence images were captured by the BioRad ChemiDoc detection system.

### Gene silencing

OCY454 cells (10^6^ cells/reaction) were transfected with the small interfering RNAs (siRNAs) ON-TARGET plus mouse Tgif1 SMART pool (Dharmacon) or ON-TARGET plus non-targeting control pool (scramble) using NEON electroporation (Invitrogen). ON-TARGET plus mouse PAK3 SMART pool siRNAs or scrambled were transfected 24 hours later to the same cells using Lipofectamine 3000 (Life Technologies). The final siRNA concentration was 70 nM/sample.

### DNA constructs and luciferase assays

The wild-type human Tgif1 promoter luciferase reporter constructs, and the Tgif1 promoter luciferase reporter construct in which the AP1 binding site was mutated have been reported previously (Saito et al., 2019). OCY454 cells were transfected with the human Tgif1 promoter luciferase reporter constructs, or the 6X-TRE-luciferase reporter (6X-TRE-luc; Clontech) to analyze AP-1 activity (Sabatakos et al., 2008), or the (−2329/149) rat PAK3 promoter, provided by Dr. Virna D. Leaner (Institute of Infectious Disease and Molecular Medicine, University of Cape Town, Cape Town, South Africa) (Holderness Parker et al., 2013), along with the humanized Renilla reporter plasmid (1:10 ratio) using Lipofectamine 3000. Luciferase assays were performed using the Dual Luciferase Reporter Gene Assay System (Promega) according to the instructions provided by the manufacturer. Firefly luciferase activity was normalized to Renilla luciferase activity.

### Promoter analysis and chromatin immunoprecipitation

Analysis of putative Tgif binding sites on the (−2329/149) rat PAK3 promoter was performed using the online platform ALGGEN-PROMO. Chromatin immunoprecipitation experiments were performed using the A/G MAGNA CHIP kit (Millipore) according to the manufacturer’s instruction. Briefly, 10*10^6^ cells of the OCY454 cell line were plated onto 15cm^2^ dishes and incubated at 37°C for 5 days. Cells were then fixed with 1% formaldehyde for 10 min, followed by 5 min of quenching using glycin. After chromatin isolation, DNA was sheared in fragments ranging between 100 and 500 bp in length through 15 cycles of high frequency sonication using a BioraptorPlus. Immunoprecipitations were performed using 7 μg of rabbit polyclonal anti-Tgif1 antibody (Santa Cruz Biotechnology) or anti-rabbit ChIP grade IgG (Abcam). The amount of pulled-down DNA was quantified by qPCR. ChIP-qPCR data were first normalized to the input of each precipitation (2 ^(Ct^ ^100%^ ^input-Ct^ ^sample)^), and then to the relative IgG sample. The gene RARα was used as positive control (Zhang et al., 2009). The oligonucleotides used to amplify the Tgif1 binding site and as negative control were the following:

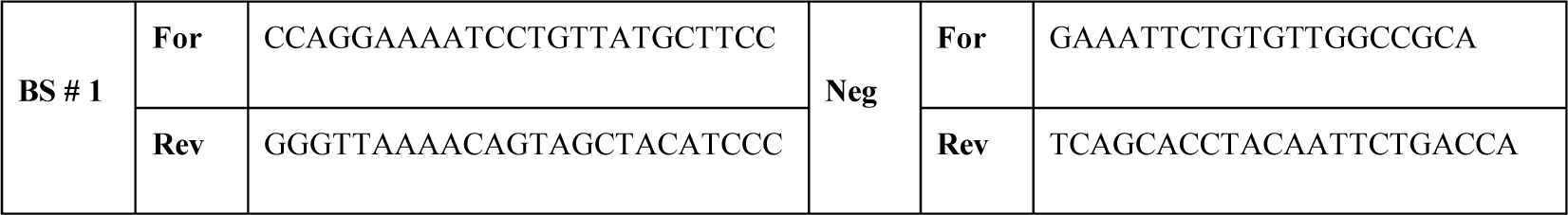

### Statistical analysis

Statistical analyses were performed using the statistical package Prism v 9.00 (GraphPad Software, San Diego, CA, USA). Statistically significant differences were determined using the unpaired T-test for comparing two groups. ANOVA followed by Bonferroni or Tuckey Multiple Comparison Test was used to compare more than two groups. The repeated measures ANOVA, estimated margins means test was used to compare the differences of two genotypes over time.

## Acknowledgements

We are grateful to A. Gasser und S. Schroeder as well as A.V. Failla and B. Zobiak of the UKE Microscopy Imaging Facility (Umif) for conceptual and technical support; G. Arndt and P. Missberger for mouse husbandry. We thank C. Walsh for providing Tgif1-deficient mice, S.J. Brandt for providing the Tgif1 reporter constructs, P. Divieti Pajevic for kindly contributing the Ocy454 cell line and V.D. Leaner for providing the rat PAK3 promoter. S. Bolamperti was supported by a postdoctoral fellowship from the European Calcified Tissue Society (ECTSABBF2015_0019). H. Saito was supported by a postdoctoral fellowship from the Japanese Society for the Promotion of Science. E. Hesse received funding from the Deutsche Forschungsgemeinschaft (HE 5208/2-1, HE 5208/2-3 and HE 5208/3-1), the AO-Foundation (S-13-73H) and the European Commission (PCIG10-GA-2011-303722). H. Taipaleenmäki received funding from the Deutsche Forschungsgemeinschaft (TA 1154/1-1).

## Supplementary Figures

**Fig. S1: (A)** Migration of calvarial osteoblasts obtained from neonatal *Tgif1^+/+^* (n=20) and *Tgif1^-/-^* (n=20) mice was analyzed using live cell imaging. The direction and distance of individual cell migration is visualized.

**(B)** Representative image of an immunoblot demonstrating the expression of Tgif1 in OCY454 cells transfected with scrambled control siRNA (Scr) or siRNA against Tgif1 (siTgif1). Actin was used as loading control.

**Fig. S2: Tgif1-deficiency does not alter the abundance or activation of the FA components and Tgif1 does not co-localize with FA complexes. (A)** Osteoblasts adhered on Col-I coated slides for 0, 20 or 60 minutes. Representative images of immunoblots demonstrating the abundance of p-FAK, FAK, p-p38, p38, p-Paxillin, Paxillin, p-src, src, Integrin β1, LRG5 and Talin. Actin was used as loading control. n=4 independent experiments. **(B)** Osteoblasts over-expressing eGFP-Tgif1 or eGFP adhered on Col-I coated plates for 60 minutes. Cells were stained for actin (magenta), nuclei (blue), Paxillin (upper panels) or Talin (lower panels, both yellow).

**Fig. S3: Lack of Tgif1 does not alter the expression of CEPs, PAK1, PAK2 or PAK4.** OCY454 cells were transfected with a siRNA targeting Tgif1 (siTgif1) or scramble control (scr) prior to adherence on Col-I coated slides for 20 & 60 minutes. Gene expression was quantified by qPCR for **(A)** Cdc42es2, **(B)** Cdc42ep1, **(C)** Cdc42ep2, **(D)** Cdc42ep4, **(E)** PAK1, **(F)** PAK2 and **(G)** PAK4. 1-way ANOVA, Tukey’s Multiple Comparison Test.

**Fig. S4: (A)** Schematic of the conserved predicted Tgif binding site on the rat and mouse PAK3 promoters. **(B)** OCY454 cells were transfected with siRNA targeting Tgif1 (siTgif1) for 48h and with siRNA targeting PAK3 (siRNA) for 24h alone and in combination alongside with the respective scrambled controls (scr) resulting in four groups: scr (scr; scr), siPAK3 (scr; siPAK3), siTgif1 (scr; siTgif1) and siPAK3+siTgif1 (siPAK3; siTgif1). Representative images of immunoblots demonstrate the abundance of PAK3 and Tgif1. Actin was used as loading control. n=3 independent experiments.

