## Supplementary figures and images for "Tgif1-deficiency impairs cytoskeletal architecture in osteoblasts by activating PAK3 signaling"

### Supplementary Figures

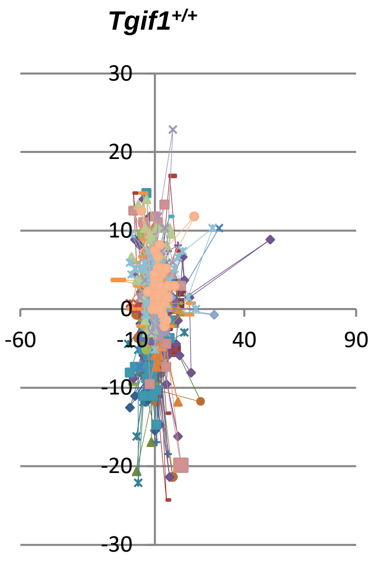

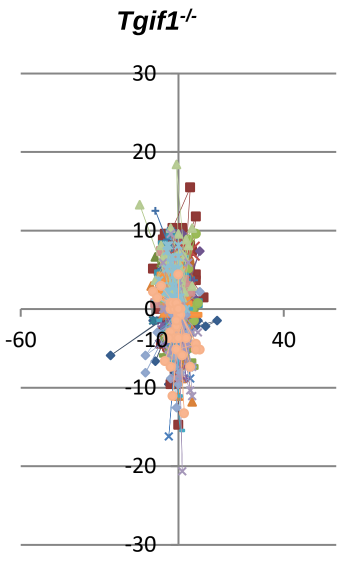

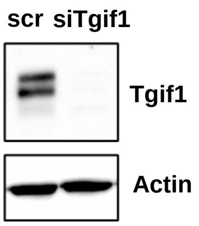

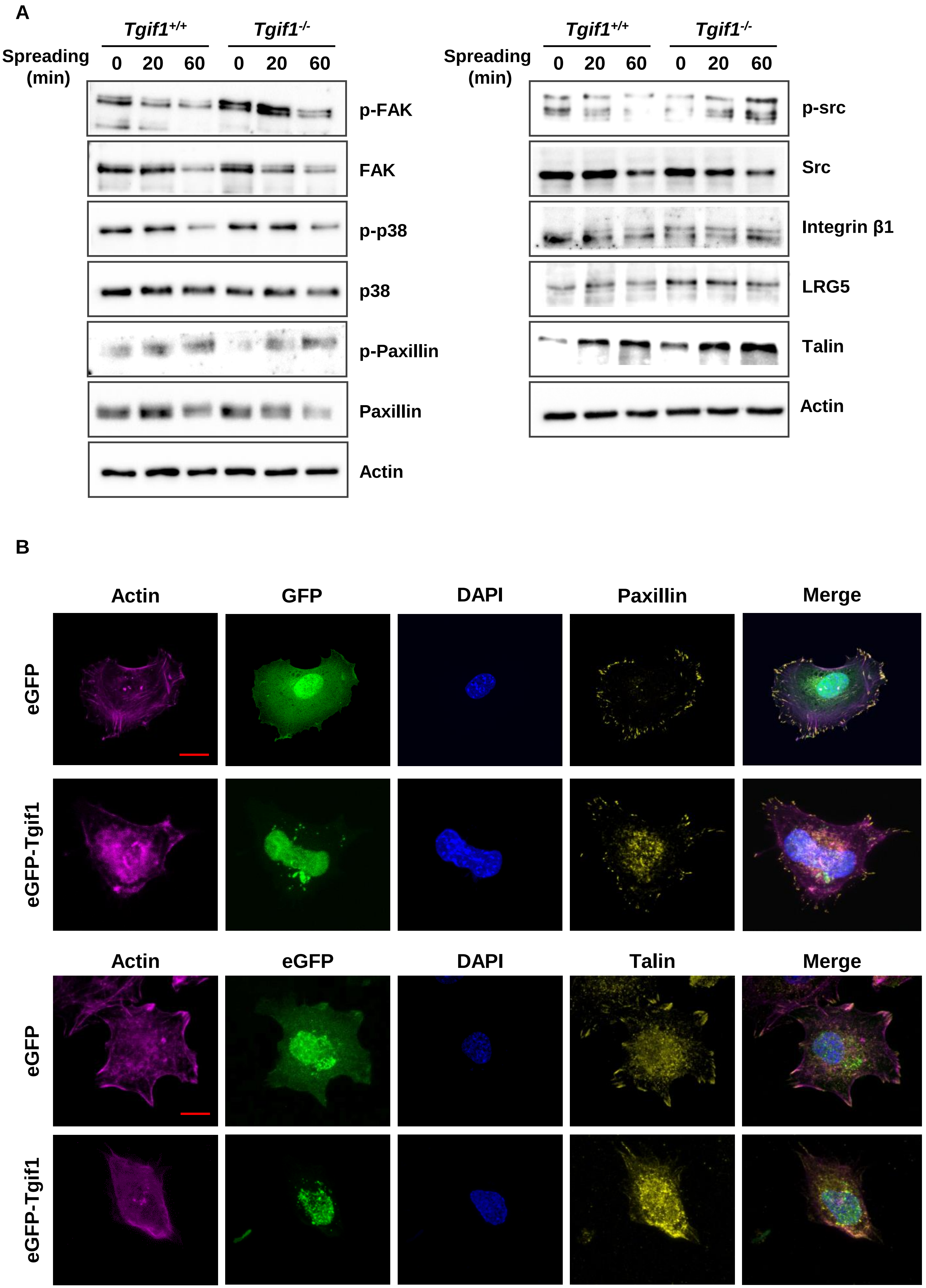

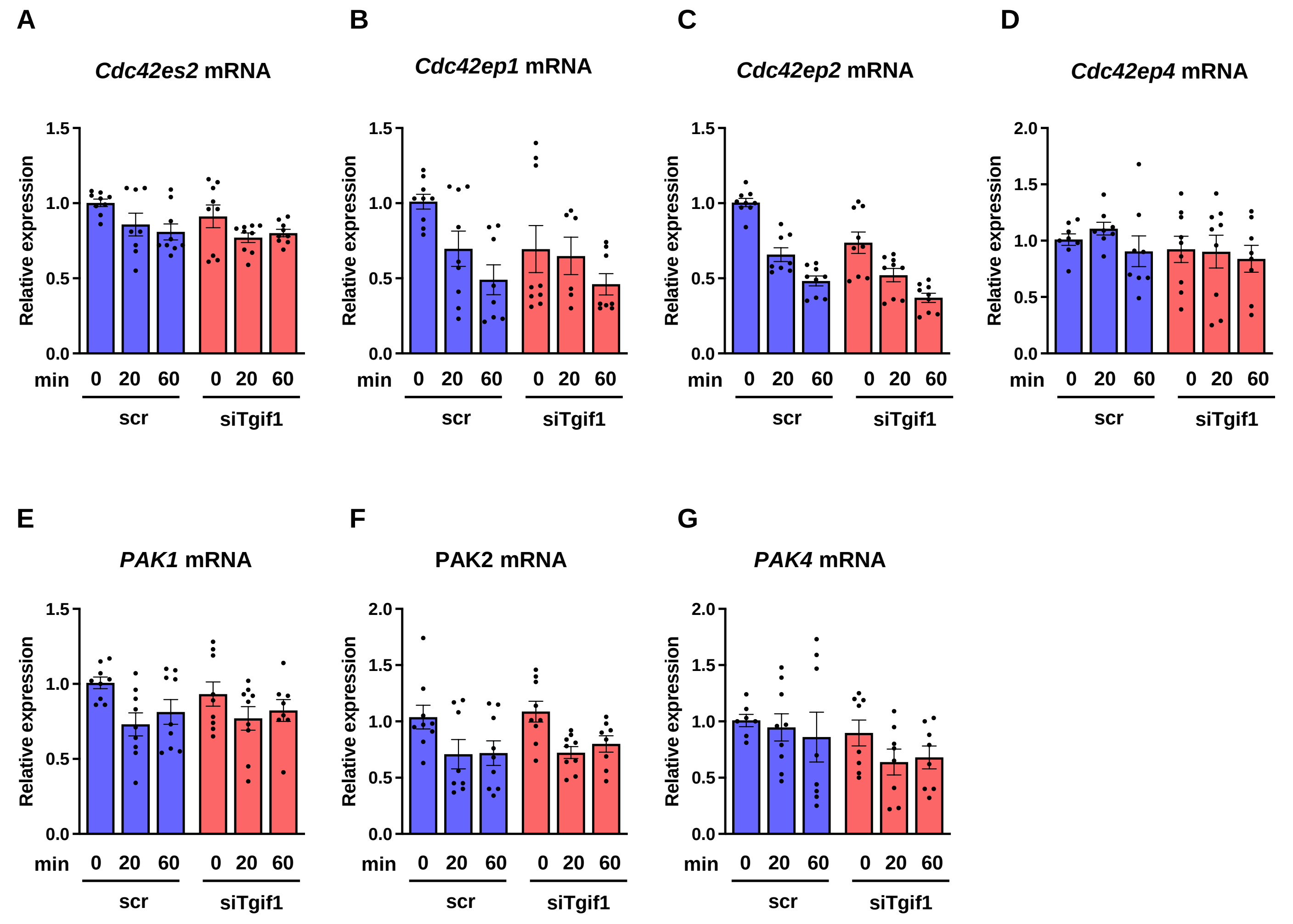

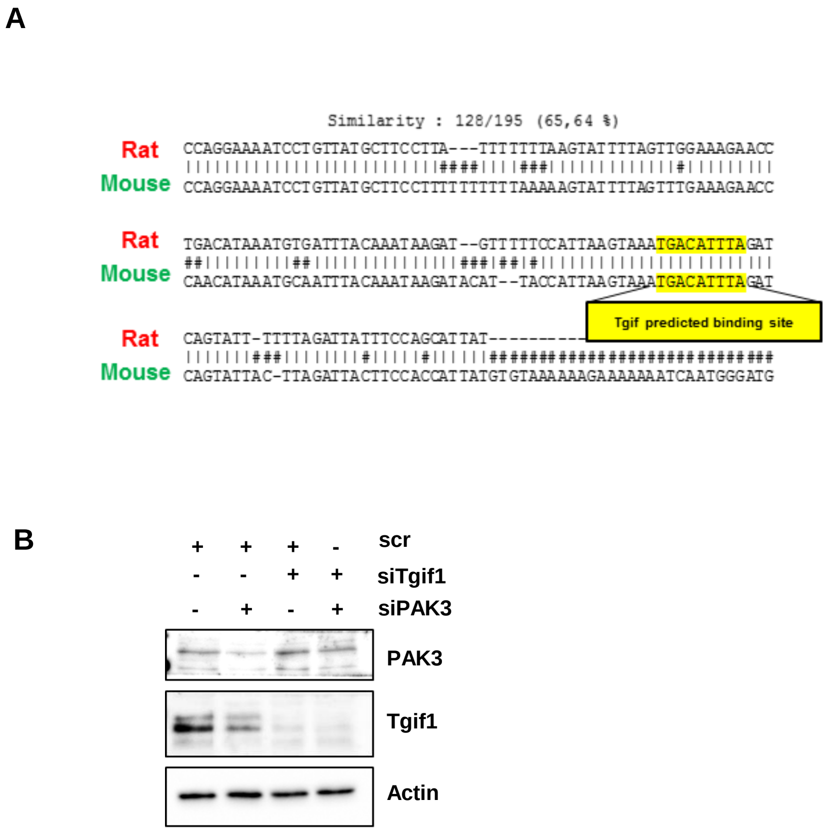
